# A mechanistic model for dynamics between CAR T cells and target cells captures features that determine killing profiles

**DOI:** 10.1101/2025.06.24.661290

**Authors:** Suriya Selvarajan, Kevin L. Scrudders, Kenneth Rodriguez-Lopez, Suilan Zheng, Bo Huang, Philip S. Low, Raghu Pasupathy, Shalini T. Low-Nam

## Abstract

Chimeric antigen receptor (CAR) T cells represent a potent, programmable therapeutic that repurposes T cell cytotoxicity toward target cell elimination. Direct killing of tumor cells has been demonstrated in several cancer contexts but deficits in destroying solid tumors have been attributed to molecular and mechanical complexity. Understanding and predicting the efficacy of CAR T cell therapy requires a rigorous framework to capture the mechanistic interactions between immune cells and tumor targets. Toward enumerating such features and to quantitatively describe tumor cell survival in response to CAR T cell visitation, we propose a parametric hazard rate model for the right-censored conditional lifetime of a tumor cell, having the form of a time-discounted integral of cell:cell engagement. The model, conditioned using data on the numbers, durations, and modes of contacts made during the stochastic encounters between the two cell types, extracts mechanistic details from the experiments. We suggest two types of parameters to encapsulate features of CAR T cell killing potency (*κ*) and the resilience of the tumor (*γ*), respectively. These parsimonious parameters, nevertheless, generate substantial insights. Firstly, phenotypic heterogeneity from interactions mediated by substrate-engaged CAR T cells dominate killing outcomes, while those initiated in suspension contribute minimally. This emphasizes that a subset of the CAR T cell population is dominating the population outcome. Secondly, the model can be used to predict the killing as a function of time upon perturbation of the system. In this case, biasing the arrival process, toward more engagement in the adherent mode, can tune the rate of killing. Collectively, insights from the model framework imply that CAR T cell accumulation and killing efficiency depend not on the sum of individual T cell interactions, but rather on the weighted, time-dependent sum of heterogeneous interactions that are environmentally modulated. The model facilitates predictive insights into how contact sequences modulate CAR T cell responses and may provide strategies to alter therapeutic regimens.

## Introduction

Chimeric antigen receptor (CAR) T cells are programmable, cell-based immunotherapies that express engineered surface receptors designed to recognize tumor-associated antigens (TAAs) and trigger destructive, native cytotoxic T cell (CTL) effector functions^1,2^. Release of lytic molecules in the intermembrane junction can trigger tumor cell apoptotic programs. Success in eliminating certain leukemias and lymphomas and the potency of single CAR T cells to kill up to dozens of tumor cells within hours^3–7^ have generated excitement in translation to the clinic. However, therapeutic application to solid tumors remains limited due to immunosuppressive and mechanical barriers in the tumor microenvironment that decrease productive cell contacts and drive diminished activation capacity of the CAR T cells^8–12^. Modeling efforts, informed by empirical data, have provided new angles on measuring the threshold criteria for CAR T cell-based elimination of tumor target cells under environmental modulation^12,13^. *In vitro* cell:cell killing assays provide access to key aspects of dynamics at the interface that control activation and destruction outcomes^14^. Observations of contacts under low effector (E) CAR T cell: target (T) cell (E:T) ratios have led to models for serial killing via submaximal degranulation of cytotoxic payloads and additive toxicity through induction of multiple, sublethal hits^5,15–18^.

Continued optimization of CAR T cell technologies have focused on maximizing signal propagation through modulating binding affinities, CAR designs, and other forms of orthogonal control^19–21^. CARs integrate an extracellular binding domain specific to a TAA with intracellular signaling domains that are expected to co-opt native T cell signaling pathways to initiate degranulation of cytotoxic molecules like perforins and granzymes^4,22^. Most TAAs are not tumor-exclusive, contributing to dangerous on-target, off-tumor toxicities^10,23^. Some of the dysfunction has been attributed to blunted CAR signaling and propagation that does not use the same machinery as the native T cell receptor (TCR)^22,24–27^. The tumor microenvironment (TME) adds additional layers of complexity through biochemical and mechanical cues that influence immune cell behavior^12,28–34^. Thus, the heterogeneous interaction landscape between CAR T cells and tumor targets remains a key challenge for therapeutic design and necessitates a deeper investigation into how mechanical and spatial features of contact influence clinical outcomes^5,9,11,35^.

Unifying probabilistic models with carefully orchestrated measurements has afforded new opportunities to infer mechanistic details from empirical data. Prior models underscore the need to extract the key criteria for successful tumor elimination and the mechanisms by which therapeutic outcomes can be bolstered^12,24,36^. These pipelines have been especially influential in capturing details that may be obscured by the inherent heterogeneity of these systems. Critical questions remain about the dynamics that control CAR T cell-mediated killing, including the density of CAR T cells required for effective tumor eradication, the determinants for recruitment of CAR T cells to individual targets, and the influence of environmental factors on these interactions^13,16,37^.

Here, we investigate and quantify the killing capacity of individual CAR T cells encountering isolated target cells in a closed, minimal environment. Understanding the outcome of CAR T cell interactions is challenging, as it involves linking the inherently stochastic behaviors of T cell scanning to tumor cell fate^38–41^. We use a second-generation CAR T cell that expresses an anti-FITC CAR that enables tumor engagement via an adaptor molecule comprised of a FITC moiety linked to a folic acid ligand^42^. Folate receptor (FOLR1) overexpression is characteristic of breast and ovarian cancers. We generated spatiotemporally-resolved data to map CAR T cell interactions with target cells to population outcomes over the first few hours after initial exposure. These data were used to develop a first-principles parametric hazard rate model that estimates time-dependent cell survival probability, conditioned on the CAR T cell visitation profile. We selected two overarching types of system parameters to reflect the efficacy of the CAR T cells (*κ*, kappa) and the resilience of the target tumor cells (*γ*; gamma), respectively. Our goal was to use minimal parameters to recapitulate the empirical observations and generate both mechanistic insights and predictive power in increasing target killing.

Two qualitative observations in the raw data were quantitatively assessed in the model. Firstly, the potency of CAR T cells that spread on the underlying, stiff substrate prior to contact with the target, called adherent interactions, are more potent. In fact, these interactions dominate the killing outcomes and those contacts that occur through CAR T cells landing on the target, termed suspension interactions, are negligible in the evolution of the system. Secondly, tumor cells are capable of increased survival likelihood following sub-lethal encounters with CAR T cells as a function of elapsed time after the encounter. Using an alternate tumor model with a slightly higher tumor-associated antigen density and an observed lower susceptibility, we show that modulation of the microenvironment primes increased CAR T cell efficacy by changing the arrival profile. The model demonstrates that characterization of a CAR T cell:tumor cell pair by the *κ* and *γ* parameters can be used to predict how to tune therapeutic outcomes and effect a target magnitude of killing within a designated time period.

## Results

### Synergy between a mechanistic model for dynamics and empirical observations

The central objective for this work was to provide a mechanistic understanding of the dynamic interactions between CAR T cells and target tumor cells (effector:target; E:T) expressed through a model. What follows is a discussion of the measurements that were used to condition the model and the considerations underpinning the model to provide the most information-rich, minimal parameter set to explain the dynamics of the system. The experiments were designed using a controlled, reconstituted system to aid in defining the fundamental criteria for productive E:T interactions. The mechanistic model was constructed to test hypotheses from the observations such as the consequences of heterogeneous interactions and the susceptibility of different tumor targets. The predictive power of the model is tested by altering the arrival process of CAR T cells to a resilient tumor target to estimate the improved killing capacity.

### CAR T cell:tumor cell interactions are marked by a phenotypic heterogeneity

We established an assay platform to screen a large number of CAR T cell:tumor cell interactions over hours and under physiological conditions. MDA-MB-231 breast cancer cells were plated on glass coverslips at single cell densities. We used a second generation, anti-FITC CAR T cell model that interacts with a tumor-associated antigen on the target in the presence of saturating concentrations of EC-17 adaptor molecules. The EC-17 ligand was designed as a small molecule adaptor of folic acid linked to a biologically inert FITC molecule in a 1:1 ratio (^43^; Fig. 1A). MDA-MB-231 cells express FOLR1 with a surface density estimated to be at least hundreds of molecules per square micron^43^. CAR T cell:target cell interactions were captured using time-lapse microscopy (Fig. 1B) using diascopic illumination over tiled fields of view covering millimeters of space and sampling at least tens of target cells. The typical temporal resolution of an experiment was three minutes which is faster than the typical time scale for killing^13,44^.

**Figure 1.**
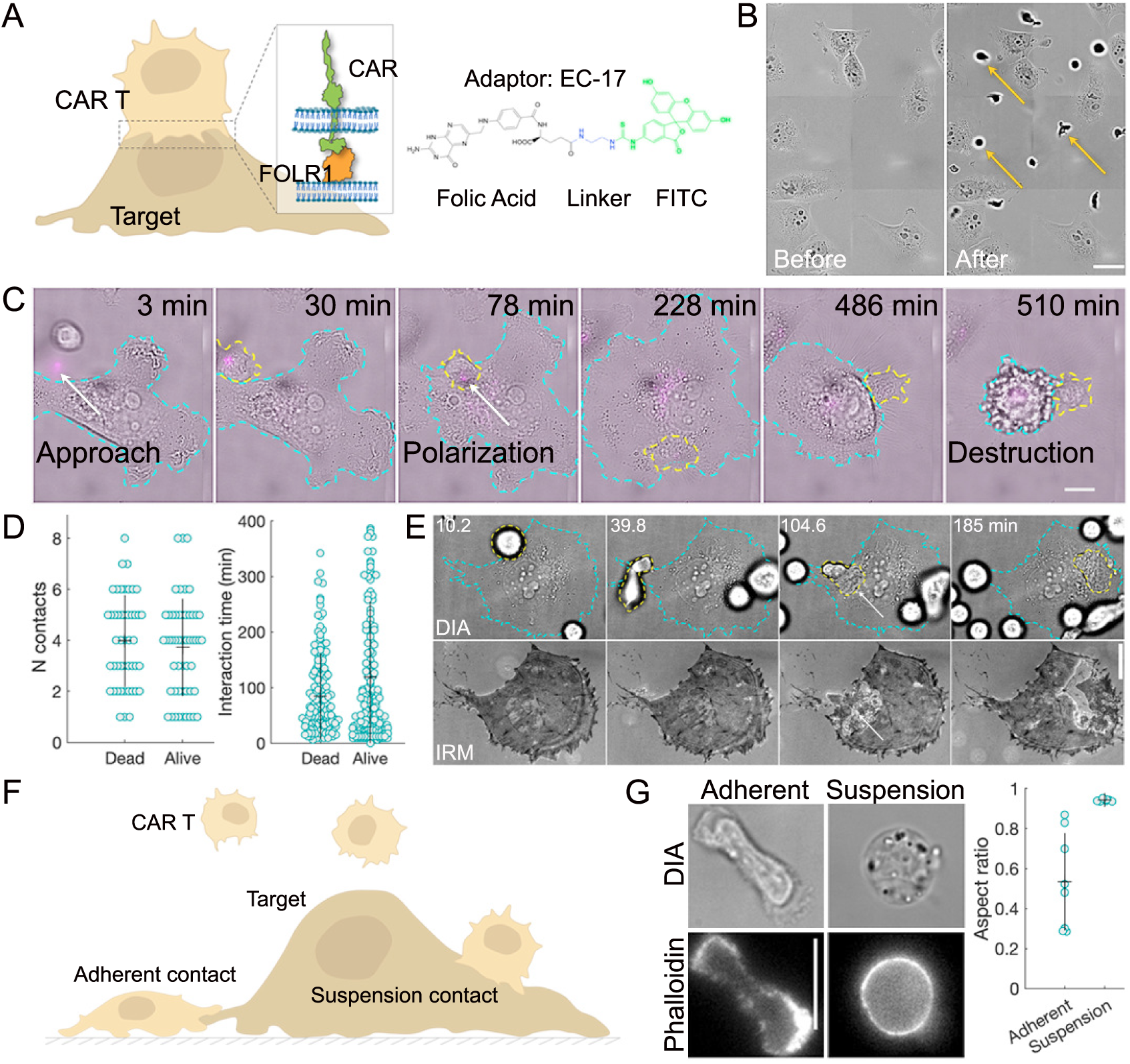
*In vitro* assay to measure features of CAR T cell elimination of tumor target cells. A. CAR T cell:target cell interactions occur at an intermembrane interface. The anti-FITC CAR binds to the FITC moiety on the EC-17 adaptor ligand that includes a linker and folic acid ligand. The latter binds FOLR1 on the tumor cell surface. B. Diascopic images of MDA-MB-231 target (T) cells plated on glass at low densities. Effector (E) CAR T cells were added (some examples indicated by yellow arrows) at a 4:1 ratio and allowed to interact for 8 hours. Scale bar = 25 microns. C. Example montage for interaction of a single CAR T cell, loaded with fluorescent LysoTracker Deep Red (shown in magenta; cell outlined in yellow) interacting with a target MDA-MB-231 cell (outlined in cyan). The CAR T cell approaches and contacts the target, polarizes its cytotoxic payload, and destroys the target. Scale bar = 10 microns. D. (Left) Single cell distributions of numbers of contacts experienced by single target cells resulting in either elimination or survival, respectively. (Right) Single cell distributions of integrated interaction times for target MDA-MB-231 cells that are either destroyed or survive, respectively. E. Montage of CAR T cell interactions with the underlying substrate in addition to those with the MDA-MB-231 target cell. Top: Diascopic imaging of a target cell (outlined in cyan) in the presence of multiple CAR T cells. One CAR T cell is outlined in yellow to indicate its position over time. Bottom: Corresponding IRM images. White arrow in each channel indicates the signature of the same CAR T cell that experiences extensive contact with the glass substrate after moving underneath the target cell. Scale bar = 10 microns. F. Schematic for two modes of contact with target cells, either in the suspension or adherent context. G. Cells in the adherent context are more anisotropic than those forming suspension contacts. Diascopic imaging (left, top) and fluorescent imaging of phalloidin staining (left, bottom) indicate differences in actin distribution in the two contact modes. Scale bar = 10 microns. Quantification of aspect ratio for CAR T cells in t5h/3e2 adherent or suspension context (right).

Effector-to-target (E:T) ratios were optimized to avoid overcrowding and ensure that each CAR T cell arrival and interaction could be quantified. Killing responses over 16 hours increased with a higher E:T ratio (Fig. S1). A ratio of 4:1 and 8 hours were used for the remaining experiments since more than half of the target cells were destroyed and, thus enabled discrimination between target elimination and survival (Fig. S1). The stages of CAR T cell interaction with a target included approach and contact and might result in polarization of the cytotoxic granule payload, its release, and subsequent death. Individual CAR T cells interacted with targets and demonstrated a range of behaviors. The shorter timescale of the measurements limited the observation of serial killing^3,5^. An example sequence is shown in which a CAR T cell loaded with LysoTracker Deep Red, to mark the low pH compartments including acidic vesicles loaded with perforins and granzymes, is shown to undergo the full activation sequence and kill a target cell (Fig. 1C and Movie S1). The collections of interactions between effector and target cells were curated based on number and duration of contacts throughout the temporal sequence. These first order parameters failed to capture the differences between target cells that were killed and those that survived Fig. 1D. The average contact time for MDA-MB-231 cells that survived was higher (Fig. 1D; dead cells: 84.4 minutes; surviving cells: 118.9 minutes), suggesting that activation decisions were made rapidly. Additionally, there were a large number of interactions that did not culminate in sufficient CAR T cell activation to contribute substantial damage to the target cells. The discrepancy between some cells that quickly committed to activation and those that suggested diminished signaling could not be explained by contacts alone.

An unexpected observation was that some CAR T cells were observed to make strong contact with the underlying glass substrate and crawl toward targets to effect contact. Occasionally, these highly spread effectors were even observed burrowing under the target cell (Fig. 1E and Movie S2). CAR T cells that engaged the substrate were distinguished from those interacting in suspension using a label-free, surface selective interference reflection microscopy (IRM) approach (Fig. 1E). Thus, we introduce two distinct interaction types: adherent contacts interacting in the high-stiffness glass environment and suspension contacts interacting from above, surrounded by fluid (Fig. 1F). All cells were detectable in the diascopic images but the additional information from the IRM channel ensured that spurious assignments of substrate contact were avoided. The remarkable differences between the two contact modes prompted us to suspect that the killing behaviors of the two interaction types were separable. A simplified prediction was that the adherent mode would be consequential and reflect the strong role for mechanics that has been observed in many immune cell-tumor cell interactions^28,29,45–47^. We note that only one target cell died exclusively through suspension interactions.

Prior evidence for substrate stiffness in priming native T cell activation implicated rearrangements in the cytoskeleton in facilitating lytic granule polarization^45,46,48^. In the two phenotypic contact modes identified here, immunofluorescent detection of actin by phalloidin labeling showed anisotropic distributions of cortical actin in CAR T cells in the adherent contact mode. These cells were also less round than those making suspension contact (Fig. 1G). Taken together, these observations are consistent with a model for enhanced killing by CAR T cells that experience high microenvironmental stiffness while contacting the target cell. Indeed, there are extensive observations that connect the mechanics of T cells and of the target to priming T cell activation and cytotoxic outcomes^46,49,50^.

### Extracting contact histories to condition mechanistic model

The interaction data were scrutinized using the concept of contact time profiles to construct a history of interactions for each individual tumor cell. Data were curated for the first 8 hours of the experiment to extract the dominant contributors that establish system-wide outcomes. An example state trace that led to target death is shown in Fig. 2A (additional examples in Fig. S2). The tumor target was alive (teal) at the beginning of the experiment. CAR T cells that interacted in the suspension (red) or adherent (blue) modes were tracked for their integrated numbers and durations as a function of time. If the target cell died (gray), the time of the onset of blebbing and apoptosis was recorded. A trace for an MDA-MB-231 cell that remained alive at the end of the 8 hours is shown in Fig. 2B (additional examples in Fig. S3). The distribution of contact times showed that the adherent-type interacts were shorter than those that occurred in the suspension mode (Fig. 2C). Analysis of the fractional time spent in each contact mode revealed several key features. Firstly, we confirmed that contacts were required for target killing. Those cells that survived spent a shorter fraction of time in contact with CAR T cells. The longer interaction times for suspension contacts led to more simultaneous binding in that mode, regardless of eventual outcome. Secondly, adherent contacts appeared to be more potent. Only 1 or 2 simultaneous contacts were necessary for the majority of the killing outcomes, whereas a large fraction of time was spent with two or more suspension contacts to generate target destruction(Fig. 2D). These data emphasized the need for a probabilistic treatment of the data to extract mechanistic information.

**Figure 2.**
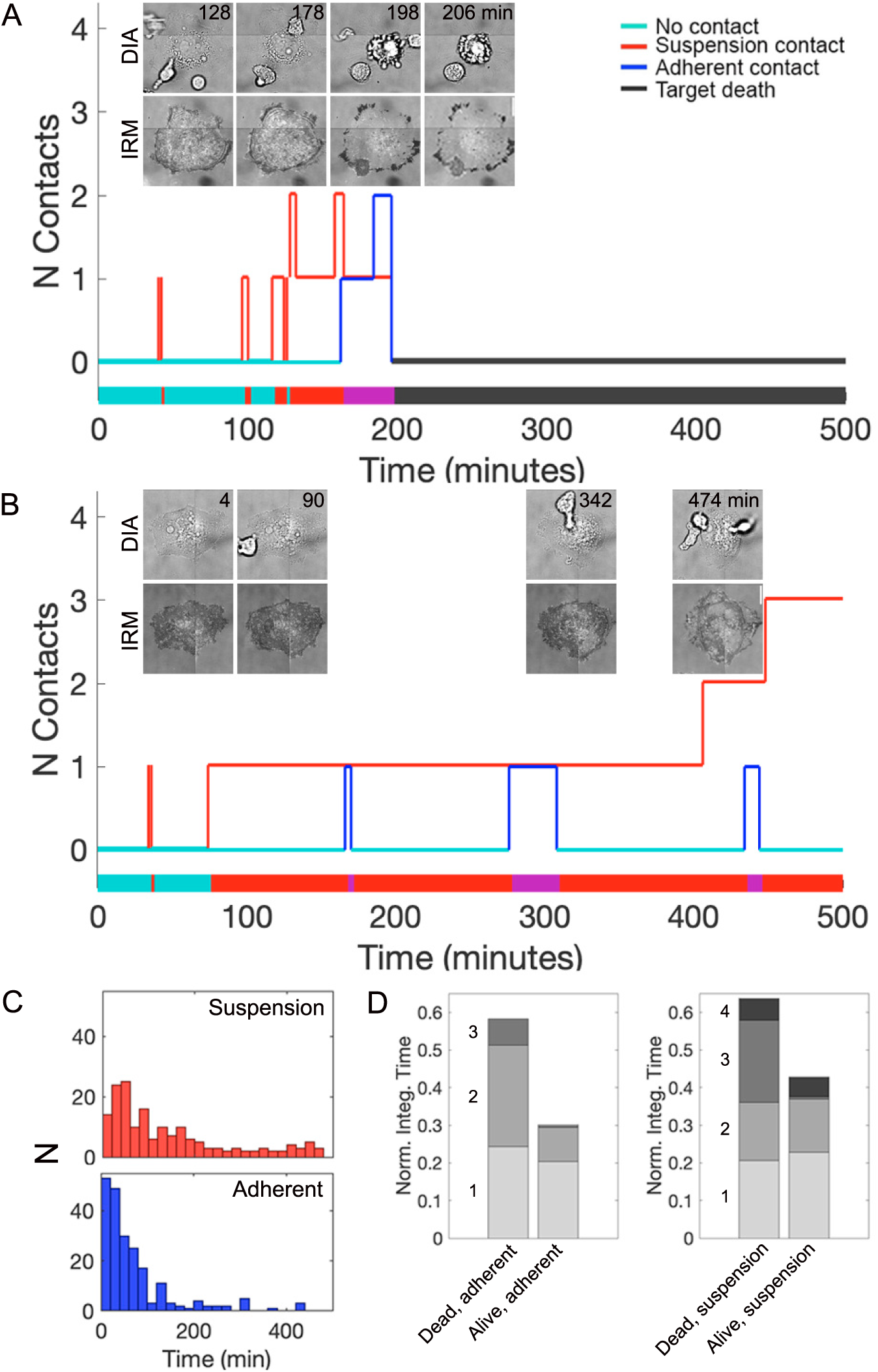
Extracting the history of contacts experienced by target cells predicted a strong contribution from adherent CAR T cell contacts to MDA-MB-231 cell death. A. Example state trace for an MDA-MB-231 cell that is destroyed after interactions with CAR T cells in the suspension (red) and adherent contexts (blue). Number of contacts are shown as a function of time. Target alive shown in teal and target dead indicated in gray. The overall history is shown as a projected timeline along the x-axis with the types of contact, where simultaneous adherent and suspension contacts are in purple, and the state of the target cell as a function of time. Insets are diascopic and IRM images of the raw data. Scale bar = 10 microns. B. Example state trace for an MDA-MB-231 cell that survives through the duration of the experiment using the same scheme as in (A). Insets are diascopic and IRM images of the raw data. Scale bar = 10 microns. C. Interaction time distributions for the suspension (top) and adherent (bottom) contact modes. D. Fraction of time spent in different contact contexts with MDA-MB-231 cells that either survive or die. Numbers of contacts from 1-4 are shown in gray in ascending order.

CAR T cell cytotoxic effects are primarily mediated through release of perforins and granzymes to induce target cell apoptosis^51^. The adherent mode could generate CAR T cell priming by promoting lytic granule polarization to the cell:cell junction^52,53^ even during brief engagements^54^. In contrast, suspension-interacting CAR T cells may exhibit less productive granzyme uptake through transient perforin pores that are rectified by the target cell^55^. Thus, mechanical activation may play a key role in induction of damage by adherent contacts. Our studies centered on quantifying the differential potency between adherent versus suspension CAR T cell interactions in driving tumor elimination, while simultaneously mapping how cancer cells responded to assault. To disentangle these dynamics, we developed a parsimonious mathematical framework using two types of parameters related to the killing capacity of the adherent and the suspension cells and the tumor repair capacity, respectively.

### A hazard rate model for CAR T cell:tumor cell dynamics

Recall that our objective was a quantitative understanding of the target’s time-dependent survival probability, upon visitation by a CAR T cell. As the mathematical means of expressing such understanding, we employed the *hazard rate*, interpreted as the instantaneous risk of the tumor cell being eliminated. Specifically, we proposed the following six-parameter model form for the time-dependent hazard rate of the tumor cell:

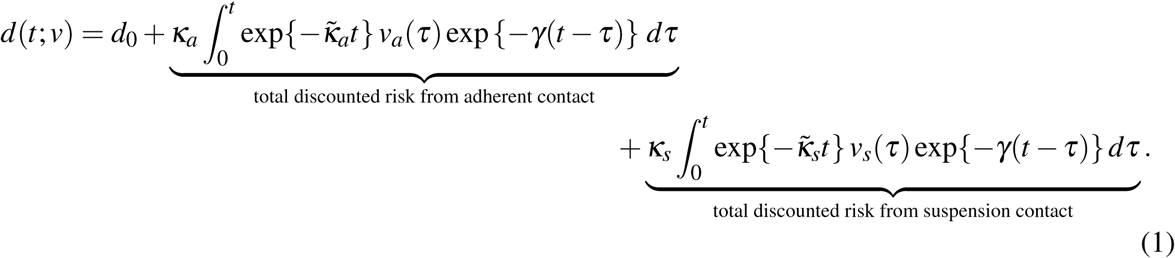

where *t* ≥ 0 represents time; *v* = (*v_a_*(*t*)*, v_s_*(*t*))*, t* ≥ 0 is the time-dependent visitation profile of CAR T cell(s), that is, *v_a_*(*t*) and *v_s_*(*t*) denote the number of adherent and suspension cells, respectively, visiting the tumor cell at time *t*; and (*d*_0_*, κ_a_*, *κ̃*_*a*_, *κ_s_*, *κ̃*_*s*_, *γ*) are the model parameters. With *d*(,) in (1) representing the hazard rate, and *L* denoting the lifetime of a tumor cell, the *survivor function* of the tumor cell can be expressed in terms of the hazard rate:

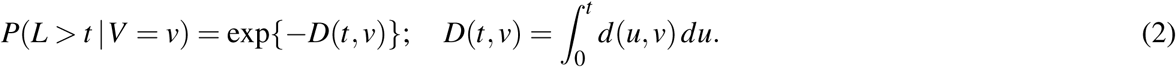

(See Methods for a full derivation of the above model.)

### Model structure and interpretation

The structure of the model in (1) was chosen for mechanistic interpretability, and to facilitate statistically testing key dynamics evident in the empirical findings, and based on hypotheses in the literature. The following observations about model structure and parameters are salient. A schematic representation of the model framework is also provided as Figure 3A.

a. The parameters *κ_a_*, *κ̃*_*a*_ are “killing coefficient” parameters specific to the adherent (‘a’) CAR T cell contact configuration, whereas *κ_s_*, *κ̃*_*s*_ are analogous parameters corresponding to suspension (‘s’) contact mode. Separate model parameters for adherent and suspension interactions allowed, simply by comparing parameter estimates along with their estimated standard errors, for statistically testing the hypothesis that substrate engagement is more potent than suspension engagement as a killing mechanism.
b. The parameters *κ̃*_*a*_ and *κ̃*_*s*_ (having the units of inverse time) were included to test the “exhaustion hypothesis,” whereby the killing potency of CAR T cells diminishes following initial stimulation or over engagement with the tumor cells. Exhaustion may be relevant to our measurements given observations of rapid onset, within hours, of the dysfunctional state^57^.
c. The damage done to the tumor cell through engagement with CAR T appears to be cumulative, and this observation is reflected through the inclusion of an integral that is weighted by the time-dependent visitation functions *v_a_*(·)*, v_s_*(·).
d. The parameter *γ* functions as a “risk discounting” parameter to reflect the observation that tumor cells seem to recover after engagement with a CAR T cell, so that encounters in the distant past were weighted less than those in the recent past.
e. The parameter *d*_0_ represents the baseline hazard rate, that is, the hazard rate for a tumor cell in the absence of any engagement with a CAR T cell.

**Figure 3.**
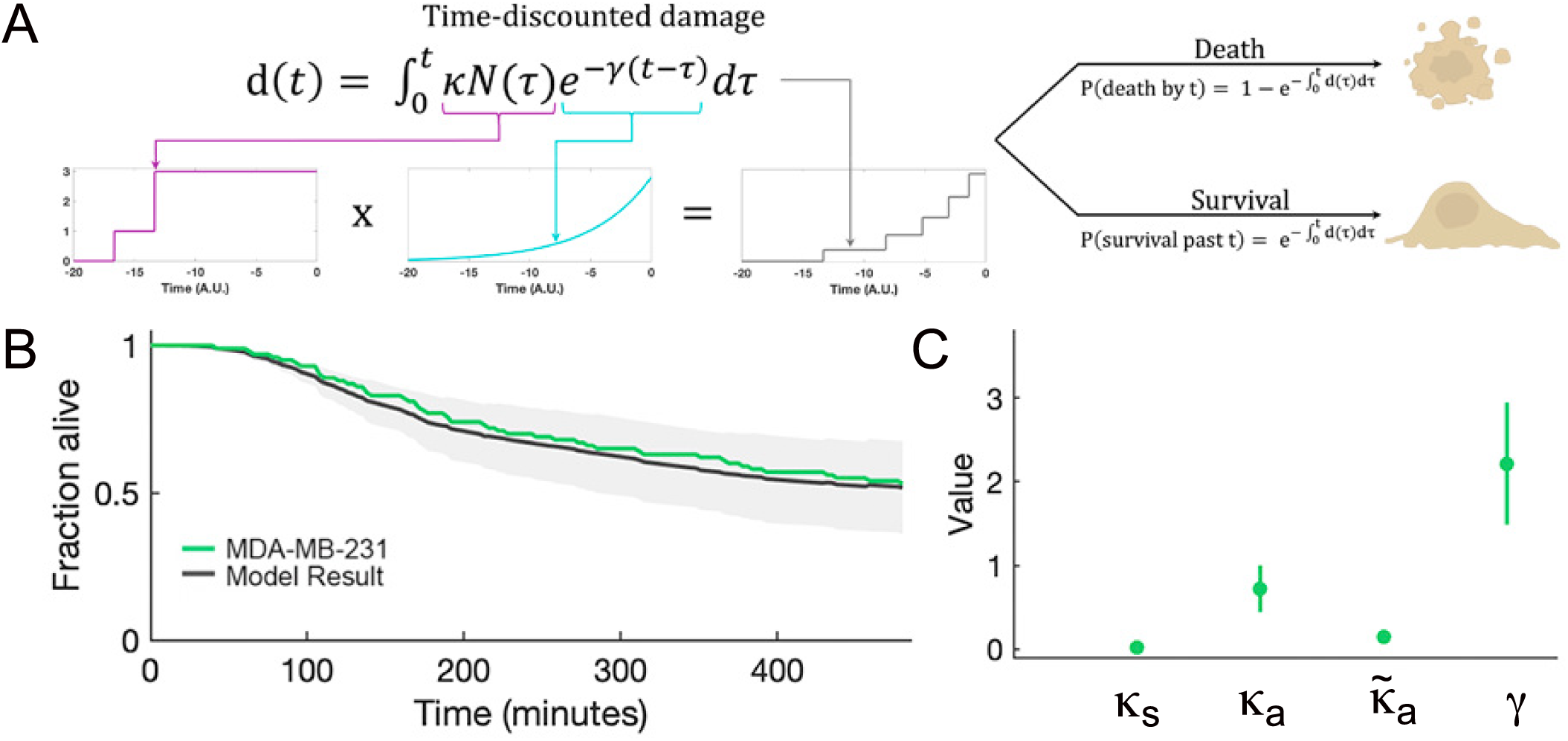
Probabilistic model using probability of killing and target recovery parameters recapitulates the killing characteristics for CAR T cell and MDA-MB-231 cell interactions. A. Schematic of the modeling framework. B. Kaplan-Meier plot for 40 simulations. The average killing curve is shown in gray with the upper and lower bound estimators shown as a light gray area around the mean. The empirical data for destruction of MDA-MB-231 target cells are plotted in green. C. Values of *κ* (adherent; a and suspension; s, respectively) and *γ* determined from the model. Values are plotted with 95% confidence interval.

### Model estimation

While the model in (1) included the complete set of parameters, the parameter estimation process evaluated which parameters were statistically different from zero and, thus, considered which parameters could inform the underlying mechanisms. For instance, suppose the estimator 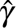 of *γ* had an estimated standard error sê(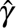) that was large compared to 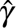. This constitutes evidence that the hypothesis of tumor recovery post engagement, might not be true. Similarly, if the estimator 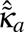 of *κ̃*_*a*_ had an estimated standard error that is small compared to its coefficient estimate, this might constitute further evidence for the CAR T exhaustion hypothesis. In general, whether or not a specific parameter in our proposed model was deemed statistically significant depended on whether the absolute value of its estimate exceeded its estimated standard error by at least a factor of two. In addition to including only parameters that were deemed statistically significant, the overall model’s accuracy was assessed by comparing the survival probability as estimated through (2) against the actual Kaplan-Meier plot, e.g., see Figure 3B. We provide further detail on parameter estimation, standard error estimation, and model selection, in the Appendix.

Another important aspect of the model in (1) pertained to how the visitation process was incorporated. Suppose again that *L* denotes the lifetime of a target, and *V* = *V* (*t*)*, t* 0 the corresponding CAR T visitation process. Then, *d*(*t, v*) in (1) represents the hazard rate of the *conditional random variable L V* = *v*, as opposed to the marginal random variable *L*. This subtle modeling choice reflected our thought that the CAR T arrival process was likely environment dependent, and should thus be separated from the cell:cell dynamics to the extent possible. Modeling the conditional hazard allows to isolate the cell:cell dynamics from that of the CAR T arrival process. Indeed, the survival curves seen in Figure 3, Figure 4, and Figure 5 were constructed conditional on the visitation of CAR T, that is, the randomness associated with the arrival process was “not integrated out” before constructing the survivor curves.

**Figure 4.**
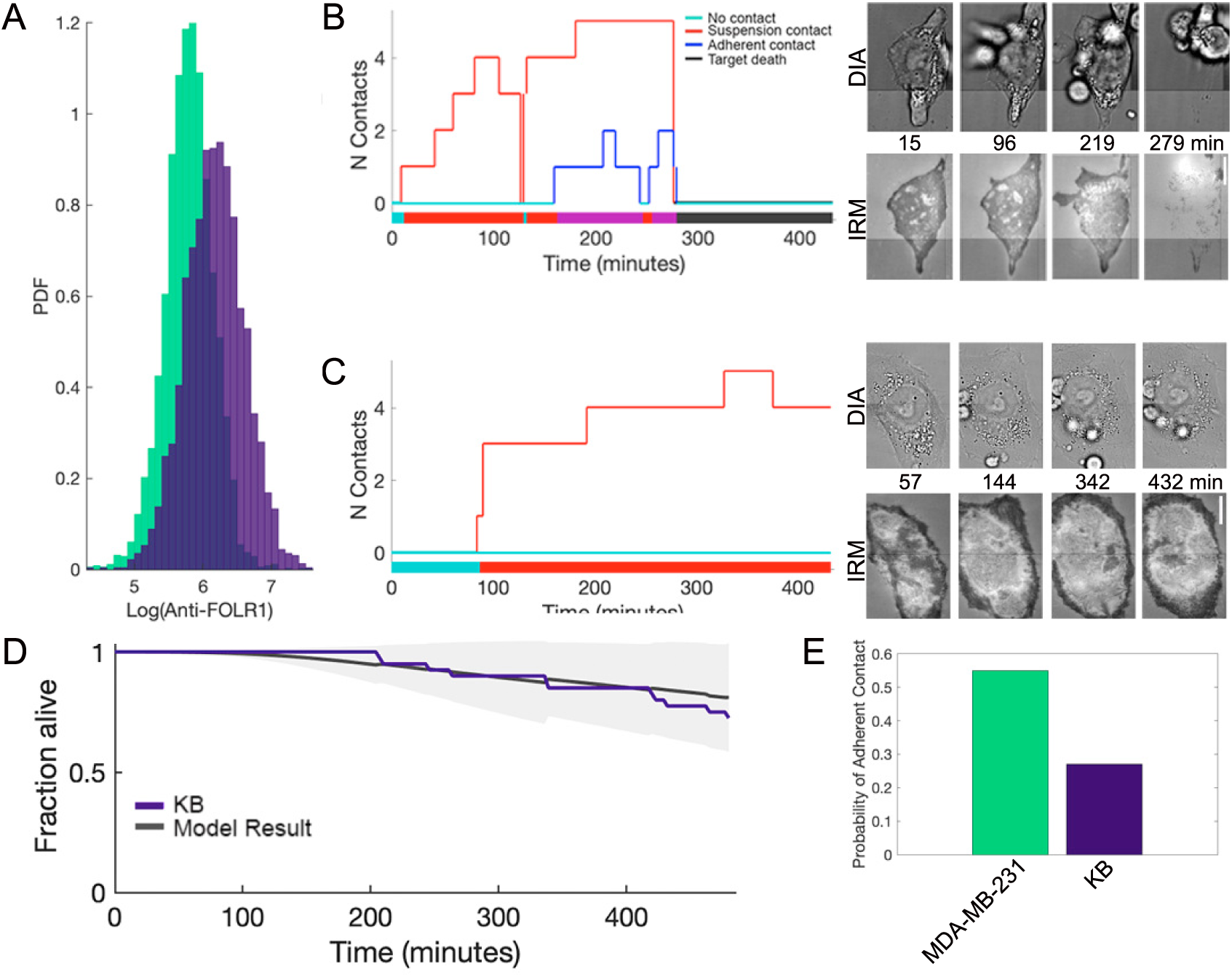
CAR T cell destruction of a different target with the same tumor-associated antigen marker. A. Flow cytometric detection of FOLR1 expression on MDA-MB-231 and KB cells. B. State trace (left) and montages in diascopic and IRM channels (right) for a KB target cell that is destroyed by interactions with CAR T cells. Montages on the right show diascopic and IRM imaging for the corresponding raw data. Scale bar is 10 microns. C. State trace (left) and montages in diascopic and IRM channels (right) for a KB target cell that is not observed to be destroyed during the course of the assay. Montages on the right show diascopic and IRM imaging for the corresponding raw data. Scale bar is 10 microns. D. Kaplan-Meier plot for 40 simulations. The average killing curve is shown in gray with the upper and lower bound estimators shown as a light gray area around the mean. The empirical data for destruction of KB target cells are plotted in purple. E. Probability of contacts in the adherent configuration for the MDA-MB-231 and KB cells, respectively. Data are combined from at least two biological replicates in each case.

**Figure 5.**
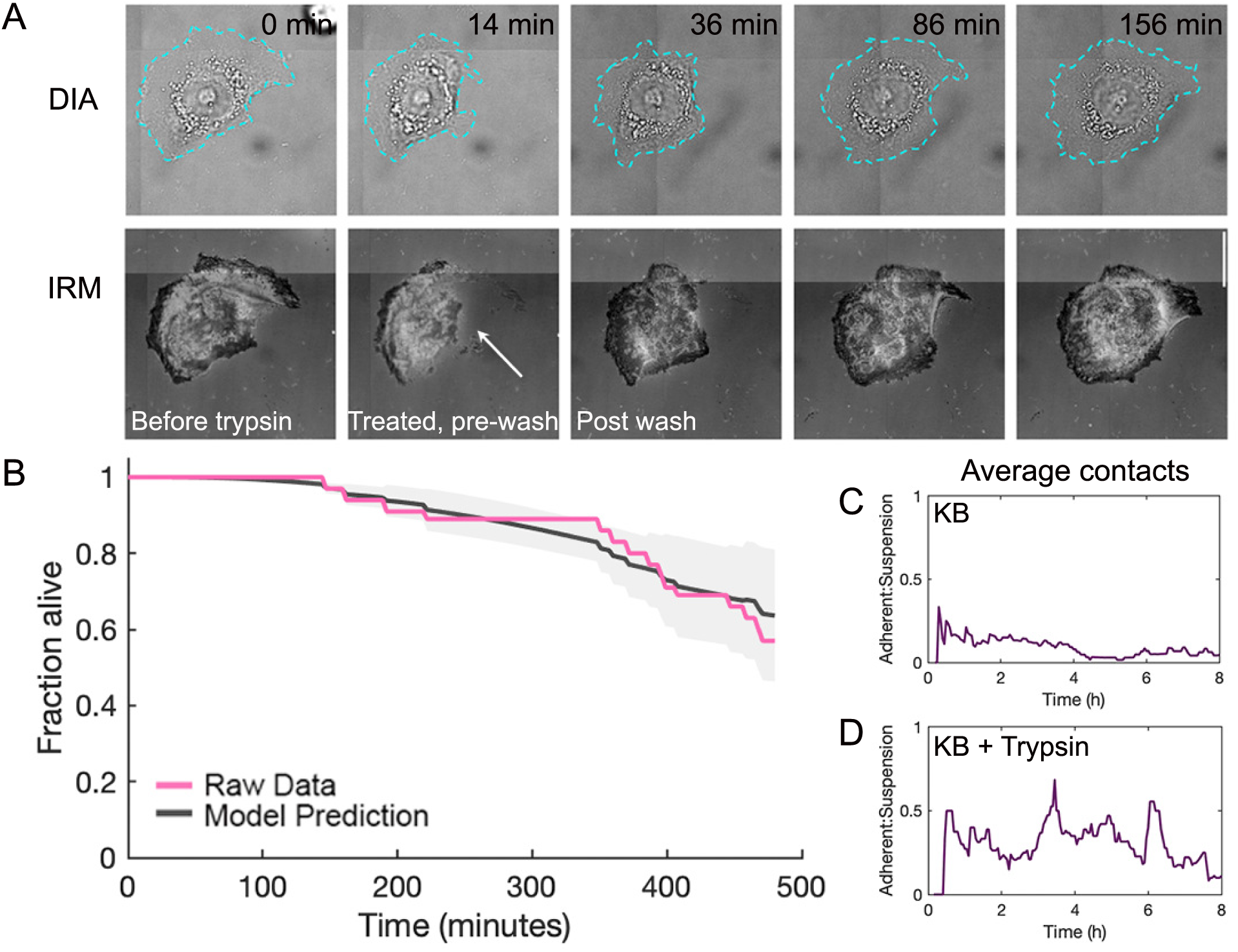
Killing outcomes are augmented by altering the arrival profile to increase adherent interactions. A. KB cell trypsinization disrupts contact with the substrate but does not kill the target cells. Montage of KB cell treated with trypsin between minutes 4-14, followed by rinse steps, and imaging in complete media. Data are shown for diascopic and IRM channels. White arrow indicates some disruption of adhesion and decrease in contact area. Respreading was detected within a few hours of the trypsinization step. Scale bar is 20 microns. Note, a slight translation in the sample occurred during the wash step, so the data have been artificially realigned to promote legibility. B. Kaplan-Meier plot for 40 simulations using the kappa and gamma values for KB cells and the visitation profile in the presence of trypsin treatment. The predicted average killing curve is shown in gray with the upper and lower bound estimators shown as a light gray area around the mean. The empirical data for destruction of KB target cells treated with trypsin are plotted in pink. C. Average number of suspension (red) and adherent (blue) contacts with KB target cells as a function of time. D. Average number of suspension (red) and adherent (blue) contacts with KB target cells treated with Trypsin as a function of time.

As we discuss further in the supplemental section, several interesting observations emerged from our model estimation process associated with MDA-MB-231. First, and as can be seen through Figure 3C, the discounting parameter *γ* is consistently significant and stable with a small standard error estimate, providing strong evidence for the presence of the tumor cell recuperation phenomenon, and the assumed discounting structure of the hazard rate. Second, both the “killing coefficients” *κ_a_* and *κ_s_* are consistently significant, and function as key explanatory variables, with *κ_a_* about a factor larger than *κ_s_*. Third, there appears to be some limited evidence of the presence of the CAR T “exhaustion hypothesis,” especially for adherent cells. Overall, the four-parameter model reported as the third entry of Table 2 seemed most favorable as measured by model parsimony, model fit, and the ability to predict the survival curve.

### The model predicts how to tune killing outcomes for a different target cell model

To probe the utility of the model framework, we selected KB cells as an alternate target that expresses the same tumor-associated antigen (TAA). The expression of the FOLR1 TAA was slightly higher than that of the MDA-MB-231 cells (Fig. 4A) which might predict that CAR T cell activation would be favored and, thus, produce a stronger killing outcome, similar to some reports^58^. Additionally, the KB cells expressed a higher surface density of the adhesion ligand, ICAM, which would be expected to favor the formation of strong cell:cell junctions and promote cytotoxic degranulation (Fig. S4). However, we observed that CAR T cell-mediated killing of KB cells was much lower than in the MDA-MB-231 case during the first eight hours (Fig. 4D). To determine whether the potency (*κ*) or susceptibility (*γ*) parameter could best explain this outcome, we applied the mechanistic model to these data. We extracted the state traces for target cells that were destroyed or that survived to condition the model (Fig. 4B-C). Like observations in the MDA-MB-231 cells, suspension binding interactions dwelled longer than adherent ones, suggesting this is a global feature of the two contact modes (Fig. S5). Most striking was the observation that the KB cells experienced dramatically fewer adherent contacts than the MDA-MB-231 cells, consistent with the weaker killing outcomes (Fig. 4D).

The smaller killing coefficient of the adherent cells and the low susceptibility of KB cells explained the poor population response. Whether the KB cells inherently disfavored adherent contacts or their microenvironment context altered the interaction profile was unclear. Thus, we predicted that modulating the arrival process in such a way as to increase adherent contacts would generate more robust destruction of KB cells. By perturbing the local microenvironment, we would phenotypically alter the behavior of microenvironment-cell interactions. The local microenvironment can profoundly impact tumor development and tune immune cell interactions^59,60^. A main barrier for cell-based immunotherapies is contact and infiltration within the tumor mass which are modulated by immunosuppressive barriers like the glycocalyx and cancer-associated fibroblasts^61,62^. The average number of contacts between CAR T cells and target cells, termed the visitation process, were extracted from the empirical observations. In the case of the MDA-MB-231 cells, the arrival of CAR T cells in the suspension and adherent configurations occurs rapidly and are present at comparable frequency (Fig. S6). KB cells treated with trypsin for a short time prior to a complete buffer wash experienced weakened interactions with the underlying substrate but retained the ability to respread and persist (Fig. 5A). This mild treatment was sufficient to alter the arrival profile of the CAR T cells and increase the frequency of contacts in the adherent configuration (Fig. 5C-D).

To test the predictive power of the model, we simulated the scenario in which the values of kappa and gamma were taken from the untreated KB cells and conditioned on the arrival profile in the presence of trypsin. Only the kappa adherent parameters were used since their predominant impacts on the systemwide outcomes were established for the MDA-MB-231 and untreated KB cells. Remarkably, the simulated killing profile matched the empirical killing behavior (Fig. 5C). Thus, the heterogeneity in contacts predominated the killing outcomes but could be modulated through modification of the arrival profile.

Table 1 summarizes the model results conditioned on the raw data for CAR T cell:target cell pair. It affords comparisons between CAR T cell efficacy and target cell resilience across the cell lines and conditions that were investigated. The values of the killing coefficient for suspension contacts are consistently negligible. MDA-MB-231 cells exhibited a high baseline susceptibility to killing by CAR T cells in the adherent contact mode (*κ̅*_*a*_ = 0.723 0.2785), although effectiveness declined moderately over time (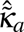 = 0.148 0.086). Notably, these cells displayed a strong recovery capacity (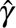 = 2.216 0.729), suggesting a strong underlying ability to recover from attacks. In contrast, KB cells showed a very low initial killing coefficient (*κ̅*_*a*_ = 0.083 0.086) and no decay in CAR T cell killing potential (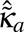*_a_* = 0.000 0.360), suggesting that CAR T cells experienced a persistently weak cytotoxic effect. Consistent with this low potency was the delayed onset of cell death; the first target was destroyed at approximately 2 hours after CAR T cell addition, which was substantially later than observed for MDA-MB-231 cells (Figure 3B and Fig. 4D). Furthermore, KB cells showed minimal recovery (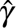 = 0.0049 0.2160), implying that while they resisted initial killing, they accumulated damage with limited resilience. Collectively, these parameters emphasize the mechanistic differences that may occur for the same CAR T cell:tumor associated antigen pair given the tumor cell genetics and environmental modulation that is present.

**Table 1.** Fitted parameters (with standard errors) for targets and conditions tested.

| Cell Type | $\hat{\kappa}_s$ (sê) | $\hat{\kappa}_a$ (sê) | $\hat{\tilde{\kappa}}_a$ (sê) | $\hat{\gamma}$ (sê) |
| --- | --- | --- | --- | --- |
| MDA MB-231 | 0.022 (0.022) | 0.723 (0.278) | 0.148 (0.086) | 2.216 (0.729) |
| KB | 0.001 (0.002) | 0.083 (0.086) | 0.000 (0.360) | 0.005 (0.216) |
| KB+ Trypsin | 0.000 (0.004) | 0.586 (0.571) | 0.501 (0.363) | 0.0000 (0.169) |

## Discussion

The molecular and physical mechanisms that regulate CAR T cell killing efficacy have been the subjects of intense investigation. Features of surface molecules, membrane topography and plasticity, and mechanical stiffness have all been implicated as regulators of cytotoxic outcomes^34,44,58,63^. Additional evidence for the collective effects of CAR T cell activation responses that generate sublethal damage suggests that complex, cooperative mechanisms are relevant in the complex, 3D tumor microenvironment^13^. In general, studies have focused on interactions shaping the intermembrane junction or the biochemical activity of the CAR T cells^31,64^. Here, we generalize the CAR T cell-dependent and tumor-specific parameters to extract the mechanistic features that are implicated in driving the system-wide killing outcomes. The model framework considers, in a simplified way, the intrinsic and extrinsic factors that impact how the two cell types respond and captures the inherent characteristics of each cell type in a single variable type. To inform the model, we combine direct measurements of cell:cell interactions and probabilistic modeling to extract the quality of those contacts. Remarkably, the parsimonious model faithfully generates time-dependent target survival traces that bear a striking resemblance to the trends measured in cell:cell killing assays.

Degranulation by T cells has been monitored through high-speed fluorescence imaging and has revealed that release of granzymes and perforins typically occurs at low levels such that only a subpopulation of granules fuse and release their payloads^44,65,66^. Submaximal degranulation of an average of 7 granules ensures that each T cell can activate, detach from the target, and continue to activate against subsequent target cells and contribute to serial killing^66^. Prior models for CAR T cell killing have emphasized that a single T cell:target cell engagement is rarely successful in generating death and implicates low levels of damage from which the target can recover^44^. Thus, a threshold level of damage is required for the elimination of the target. Our data argue that the threshold criteria for target death is not defined by a setpoint of accumulated damage but results from a distribution of heterogeneous interactions whose contact mode impacts the integral interaction time necessary for target cell death. Mechanistically, our model emphasizes that the shaping of the CAR T cell:tumor cell interface, especially mechanically, is more highly deterministic.

Extensive studies have linked physiochemical parameters of T cell:target cell interactions to killing outcomes. Many studies suggest that CAR T cell killing efficacy scales with TAA surface density and E:T binding duration^44,58,67^. In contrast, we show that the more brief adherent-type interactions were most consequential and that increased FOLR1 expression on KB cells was associated with more poor killing outcomes during the first 8 hours following exposure to CAR T cells. By investigating highly isolated target cells, we measured threshold criteria for activation and target destruction that would be expected to be more convoluted in the contexts of target cells experiencing contact and in 3D. However, we predict that heterogeneous contacts that have mechanically-driven features like the adherent mode may be distinguishable from other kinds of interactions with lower microenvironmental engagement. Thus, understanding that the suspension contacts, as described here, are inconsequential highlights the need to increase the sampling of differential contacts in tumors. To promote the discrimination of these kinds of complex data, advances in the application of artificial intelligence to segment data and parse contacts from the effector or target side will be informative^68^. This was recently demonstrated in NK cell:tumor cell interactions and demonstrates the application to immune cell:tumor cell data, in addition to setting the stage for mechanistic comparisons between different immune cell degranulation behaviors^68^.

Increased matrix stiffness, often observed in solid tumors due to extracellular matrix remodeling, can impair T cell motility, infiltration, and activation, thereby reducing their cytotoxic effectiveness. Studies suggest that stiff tumor microenvironments contribute to T cell exhaustion and dysfunction by altering mechanotransduction pathways and integrin signaling, leading to a diminished killing capacity^34^. However, some evidence indicates that moderate stiffness can enhance T cell activation by promoting immune synapse formation and mechanosensitive signaling^29^. These findings highlight the complexity of mechanotransduction in T cell-mediated tumor eradication. The probability of successful killing of tumor cells can be explained by the killing coefficient of the adherent cells. Adherent cells are consistently more potent, a trend observed in both cell lines tested. Thus, the opportunity to bias the visitation process toward the adherent contact mode is expected to generate stronger killing outcomes. Interestingly, trypsin-treated KB cells demonstrated a dramatic increase in susceptibility to CAR T cell killing (*κ̅*_*a*_ = 0.586 0.571), likely due to microenvironmental changes that enhancd CAR T cell access or activity. However, the increased sensitivity was accompanied by a sharp rise in killing decay rate (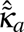 = 0.501 0.363), indicating a rapid decline in CAR T cell effectiveness over time. The KB cell recovery remained negligible (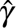 = 0.000 0.169), suggesting a compromised ability to repair damage. Collectively, changes that create a more permissive environment for the adherent contact mode is need for CAR T cells to mount an effective attack.

Therapeutic application of CAR T cells typically uses a dosing strategy that is either based on the weight of the patient with regimens on the order of 1 million CAR T cells per kilogram or a flat dose of numbers of cells, often in the billions of cells per adult patient. In response to mechanistic insights that point to environmental modulation of immunotherapies, some therapeutics have been designed to modify the microenvironment to promote increased infiltration, decreased camouflage by tumor targets, and depletion of other cell types that regulate CAR T cell killing outcomes. Our findings that CAR T cell activities are heterogeneous and there is not a correspondence between TAA expression and sensitivity point to adjusting dosing strategies for CAR T cell patients. We show that strategies to modify the local microenvironment in a way to alter the visitation process and favor the adherent contact mode result in enhanced killing outcomes. More importantly, our model enables the prediction of the extent of killing over a given time course. The unification of a model framework and characterization of CAR T cell potency and target cell resilience may provide a next step toward personalized immunotherapies.

## Methods

### Reagents

LysoTracker^TM^ Deep Red (Thermo) was used to load low pH vesicles in CAR T cells for polarization experiments. The adaptor molecule EC-17 (MedChemExpress) was a conjugate of folic acid and FITC.

### Cell Culture

MDA-MD-231 cells and KB cells were cultured in phenol free, high glucose DMEM (Gibco) supplemented with 10% (v/v) fetal bovine serum (FBS), 1% (v/v) penicillin/streptomycin (Gibco), 2 mM L-glutamine, and 1 mM sodium pyruvate and RPMI 1640 medium (Gibco) supplemented with 10% (v/v) fetal bovine serum (FBS), 1% (v/v) penicillin/streptomycin, and 2 mM L-glutamine, respectively. Primary CAR T cells expressing the anti-FITC CAR were provided by the Low lab. Briefly, T cells were isolated from healthy donors a purified by immunomagnetic selection. 293T cells were cotransfected with the CAR-encoding epHIV7 lentiviral vector and packaging vectors for lentiviral production. T cells were spinfected with lentiviral particles and expanded prior to cryopreservation ^43^. CAR T cells were cultured in TexMACs GMP phenol-free medium supplemented with 100 I.U./mL IL-2 and 2% human serum. Cells were maintained at 1-2E6 cells/mL. Cell cultures were maintained at 37 ^◦^C and 5% CO_2_.

### Microscopy setup

Diascopic and interference reflection microscopy (IRM) imaging were performed on a Nikon Eclipse Ti2 inverted microscope equipped with a motorized Perfect Focus system and motorized stage. Imaging was conducted using a 100x oil TIRF objective (NA 1.49). Illumination for epifluorescence imaging of CAR T cells labeled with LysoTracker was provided by an LED (X-Cite 110LED) lamp. Images were captured using an EM-CCD camera (iXon Life, Andor Inc.). Exposure times, multidimensional acquisitions, and time-lapse imaging were controlled using Nikon Elements software.

### Live Cell Imaging

Live cell imaging buffer (LCB: 1 mM CaCl_2_, 2 mM MgCl_2_, 20 mM Hepes, 137 mM NaCl, 5 mM KCl, 0.7 mM Na_2_HPO_4_, 6 mM D-glucose) was prepared and supplemented and 1% (w/v) BSA just prior to use. MDA-MB-231 or KB cells (50,000 cells) were plated onto sterile No. 2, 40-mm diameter round and allowed to spread for at least 10 hours in an incubator. To prepare cells for imaging, the coverslip was transferred aseptically to an Attofluor^TM^ Cell Chamber (Thermo) and equilibrated in LCB. The chamber was mounted on the microscope in a Tokai Hit stage top incubator and allowed to equilibrate to 37 ^◦^C and 5% CO_2_. Physiological conditions were maintained throughout the experiment. Time zero for a cell:cell killing experiment was designated as the time at which EC-17, at a final concentration of 1000 µM, and CAR T cells, at the desired effector:target (E:T) ratio were simultaneously added to the chamber, in suspension. Imaging was performed using DIA and IRM channels for 8 hours, with a time resolution of 3 minutes. The cell:cell killing assays with MDA-MB-231 and KB target cells included 3 biological replicates and the KB+trypsin condition included 2 biological replicates.

For imaging with loaded vesicles, CAR T cells were spun down (300xg, 4 min., at 4 ^◦^C) and the culture medium was exchanged for 1x PBS containing 100 nM LysoTracker Deep Red. Cells cell incubated at 37 ^◦^C and 5% CO_2_ for 30 minutes and then washed 3 times with LCB. Cells were kept in an incubator and used within 2 hours of labeling.

### Adhesion disruption assay

Following the normal plating of KB cells on a glass coverslip and mounting in the imaging chamber, cells were treated for 10 minutes with 1% trypsin (Gibco) followed by three rinses to fully exchange to LCB. Similar to all other cell:cell killing assays, CAR T cells and EC-17 were added to begin the measurements. The assay was run at a 4:1 E:T ratio.

### Immunofluorescence

CAR T cells were allowed to interact with MDA-MB-231 cells for one hour. Cells were then fixed at room temperature for one minute using a 0.5% paraformaldehyde (PFA; EM Sciences) solution in PBS. Cells were washed three times with 1x PBS. Following fixation, the cells were permeabilized for three minutes using a 0.2% Triton X-100 solution in PBS at room temperature, followed by another round of washing steps. A blocking step using 1% (w/v) solution of bovine serum albumin (BSA) in PBS was performed for 15 minutes. Phalloidin A555 (Thermo) was diluted in the BSA solution and applied to the cells. Following staining, the cells were washed three times with 1x PBS. To visualize F-actin localization, epifluorescence microscopy was performed using the 560 nm channel.

### Flow cytometry

Adherent cells were trypsinized to remove them from the tissue culture plastic, centrifuged, and suspended in 1x PBS at a concentration of 1E6 cells per milliliter. Labeling was performed on ice for 1 hour using Alexa-488-conjugated antibodies recognizing either FOLR1 (R&D Systems; FAB5646G) or ICAM (Novus; MEM-111). Measurements were made on a BD Accuri 6 Flow Cytometer using 488 nm excitation. At least 2000 cells were evaluated after gating unlabeled cells.

### Image Analysis

To analyze actin distributions, an analysis pipeline was to measure phalloidin distributions at the cell periphery. The perimeter of the cell was manually generated in Fiji is just ImageJ (FIJI). Line profile intensity measurements were obtained along the boundary of individual cells. These profiles were used to calculate the coefficient of variation (CV) of pixel intensities, providing a quantitative measure of cortical actin distribution heterogeneity at the cell edge. This analysis was carried out for 9 adherent and 9 suspension cells from 2 separate experiments. To further characterize cell morphology, we extracted standard shape descriptors from FIJI, Aspect Ratio (AR). The morphological feature were used to compare the two cell types. Statistical analysis and data visualization were performed using the scipy.stats and pandas libraries in Python.

### Data extraction

The data is extracted and represented as a set of observations, where each key corresponds to a unique cancer cell, and the associated values represent various events. The car t cells are manually tracked to make a recording with each cancer cell treated independently. Each event is recorded with the start time and end time of contact intervals, along with a categorical value that possibly indicates the type of contact (adherent or suspension). The data is stored in a Python dictionary, and the corresponding code for extraction is provided. The history of contacts is extracted and plotted using standard Python libraries, as shown in Figure 3. Plotting was done using MATLAB (MathWorks).

### A Mechanistic Model for Target Cell Death due to CAR T Interactions

Recall that it was our interest to construct a “first-principles” mechanistic model that explains the survival behavior of a tumor cell subject to visitation by CAR T cells. In the service of such modeling, suppose that the tumor cell is observed over a horizon [0*, h*], and suppose *L* ∊ ℝ_++_ denotes the (potentially unobserved) time of death of the cancer cell, and let *T* represent its (observed) right-censored counterpart, that is, the right-censored lifetime (RCL) of the cancer cell. Let *Y* 1, 0 indicate whether the tumor cell perishes during the fixed horizon [0*, h*]. To ease exposition, we use the ensuing subsection to first derive the density function for (*T,Y*) in terms of the hazard function *λ_L_*(.) and the survivor function 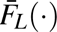 associated with *L*.

The derivation of the density of (*T,Y*) is a “marginal derivation” in that no visitation information of CAR T is used. To incorporate this key information into the model, in the second subsection that follows, we hypothesize a parametric model for the hazard function of the *conditional random variable L V* = *v*, where *V* denotes the visitation process of the CAR T cells. The hypothesized model is then used to derive the density function of the conditional random variable (*Y, T*) *V* = *v*, which in turn leads to an expression for the log-likelihood and a path to estimate the model parameters through optimization.

### The RCL Density in Terms of the Hazard and Survivor Functions

Recall that [0*, h*] denotes the time horizon of observation, that *Y_i_* = 1 if the *i*-th tumor cell survives during this period, and *Y_i_* = 0, otherwise. Furthermore, let *L*_1_*, L*_2_*, …, L_n_* denote independent and identically distributed (iid) random variables corresponding to the times at which the tumor cells perish in the hypothetical situation where the horizon is observed ad infinitum. It is important that the times *L_i_, i* = 1, 2*, …, n* are right-censored, in that *L_i_* cannot be observed past the horizon [0*, h*]. Denote the random variable corresponding to the “observed” times

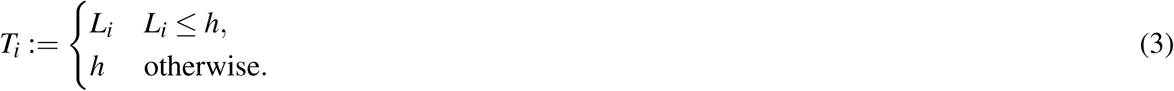

If the lifetimes *L_i_, i* = 1, 2, …, *n* have probability density function (pdf) *f_L_*, the random variable 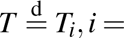 1, 2, …, *n*, and the random variable 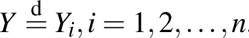, then it is seen that the joint pdf of (*T,Y*) is

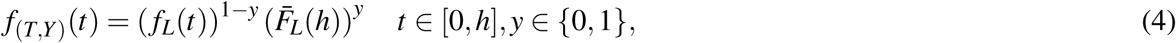

where the *survivor function*

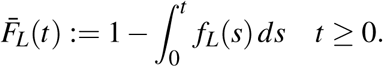

Our main modeling instrument will be the *hazard rate* [69, p. 608]:

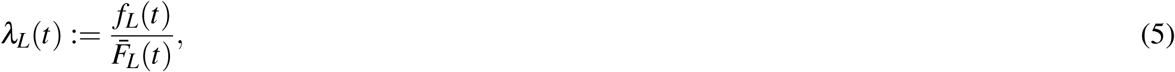

and the *hazard function* [69, p. 608]:

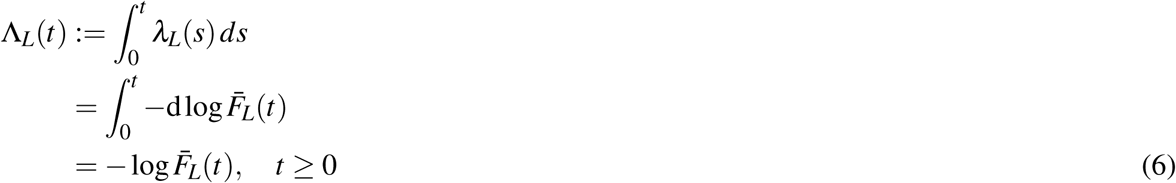

associated with the lifetime random variable *L*. The expression in (6) can be used to construct the expressions for the survivor function and the lifetime pdf, expressed in terms of the hazard rate and the hazard functions:

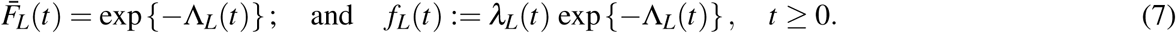

Plugging the expressions obtained in (7) into the joint pdf derived in (4), we get an expression for the joint pdf of (*T,Y*) in terms of the hazard rate and hazard function:

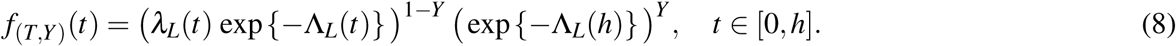

As a broader commentary that may be relevant for all modeling endeavors in similar contexts, we list some further observations. The expression appearing in (1) is a parametric model for the *hazard function*, as opposed to a *survivor function* or a *density function*. It is well-known in reliability^39^ and survival analysis^41^, that the hazard function is a more intuitive modeling object due to its interpretation as the instantaneous risk of an event happening. Our past and current experience corroborates this broader modeling insight — it is easier to posit a model on the instantaneous risk of the target cell being neutralized, as opposed to the likelihood of the target surviving for some specified length of time.

### The RCL Density Conditional on Visitation

Let *V_i_*(*τ*) = (*V_a,i_*(*τ*)*,V_s,i_*(*τ*))*, τ, i* = 1, 2*, …, n* denote the iid *visitation process* to the *i*-th tumor cell. The process *V_i_* = (*V_a_,V_s_*) is a vector process consisting of the adherent and surface visitation processes. Then, we hypothesize the following parametric model for the hazard rate of the conditional random variable *L* |*V* = *v*, where 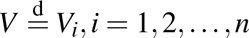:

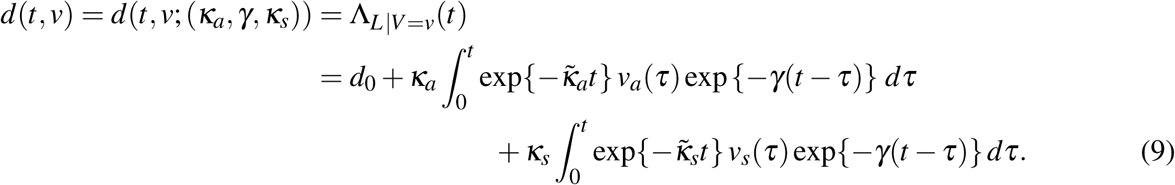

We also obtain the corresponding hazard function of the conditional random variable :

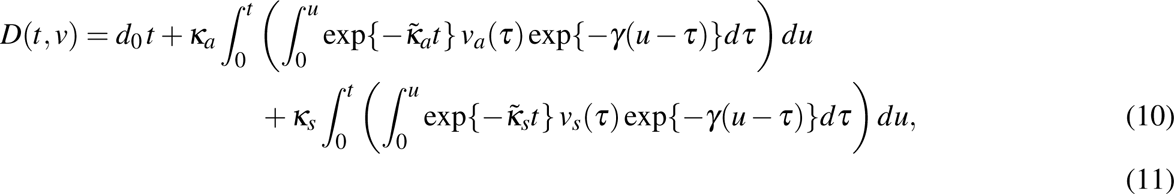

which by definition is 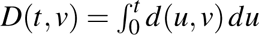. Using (9) and (10), and retracing the steps leading to (8), we obtain the pdf of the conditional random variable (*T,Y*) |*V* = *v*:

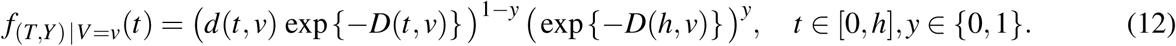

The derivation of the joint density of the condition random variable *L|V* = *v* (as appearing in (12)) represents the key step in the modeling process. For instance, as we discuss in a subsequent section on likelihood calculation, the density in (12) when combined with observed data on cell death times and the CAR T visitation process, provides a standard route [70, Chapter 6] to estimate the embedded parameters (*d*_0_*, κ_a_, κ_s_, γ*) in the model. Once such estimation is accomplished, objects of interest, e.g., the survivor function of the cancer cells, can be estimated.

### Likelihood Calculation and Inference

Given observations (*T_i_,Y_i_, v_i_*)*, i* = 1, 2*, …, n*, the (conditional) likelihood function based on the derivation of the previous section follows in a straightforward manner:

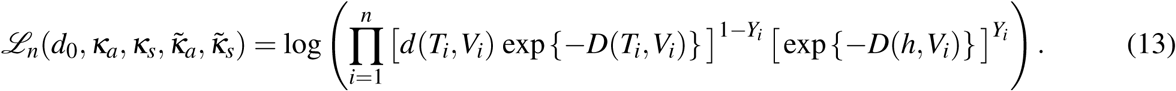

From the expression in (13), we can obtain an estimator 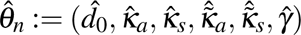 of the parameters θ_0_: = 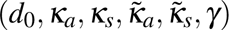 by maximizing the log-likelihood function *L_n_*(.) with respect to 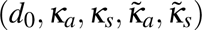, that is, by solving the following optimization problem:

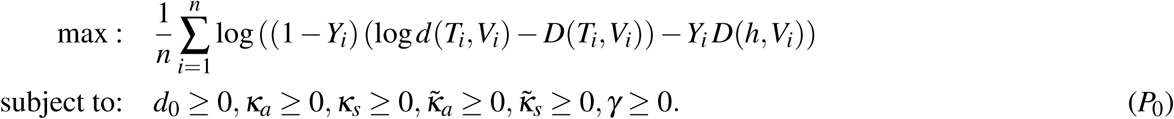

We estimated the standard errors associated with the solution 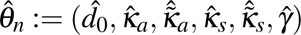 of (*P*_0_) through an asymptotic approximation. Specifically, assuming that the function

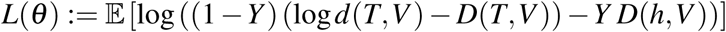

attains its minimum at *θ* = *θ*_0_, it is well-known that under fairly mild conditions^71^, the maximum likelihood estimator 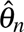 of *θ*_0_ exhibits a central limit theorem of the form

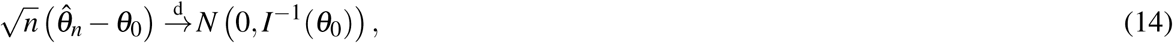

where *I*(θ_0_) = − ∇^2^*L*(θ_0_) is called the Fisher information matrix. The CLT in (14) suggests approximating the standard error of 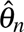 as

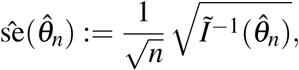

where the matrix 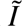 is obtained using an appropriate finite-difference approximation of *L*, and the square root operation on the matrix is allowed since we assume *I*^−1^(θ_0_) is positive definite.

## Supplemental Information

### Model Estimation

Recall that we modeled the hazard rate associated with the tumor’s conditional lifetime using the function

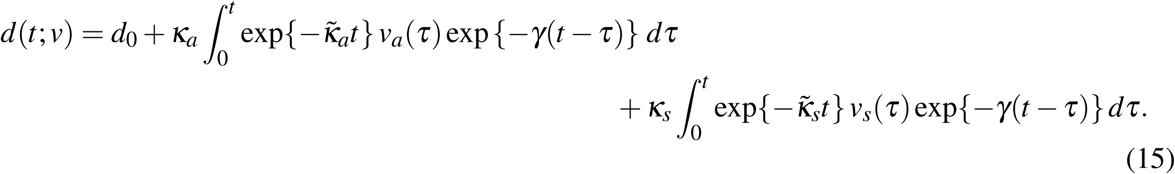

In (15), *t* ≥ 0 represents time, *v* = (*v_a_*(*t*)*, v_s_*(*t*))*, t* 0 is the time-dependent visitation profile of CAR T cell(s), and 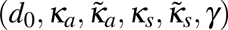 are the six model parameters. Parameter estimation using maximum likelihood, as described in the section on methods, was accomplished by numerically solving the optimization problem in (*P*_0_). Although solving (*P*_0_) is non-trivial in general, the small number of decision variables allows for an exhaustive grid search. Our numerical algorithm exhaustively explores the parameter space by iterating over a small pre-specified discretization for each element of the parameter vector 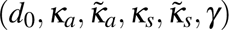.

Table 2 that follows lists estimation output from ten of the most interesting models that were estimated using the optimization procedure just described. The first six columns in Table 2 list parameter estimates, along with standard error estimates listed within parenthesis. (All parameter estimates and standard error estimates are denoted with a “hat,” since these are estimates that will tend to their true counterparts only as the number of observations in the dataset tends to infinity.) An “NA” for the parameter estimate indicates that the particular parameter was deliberately excluded from the model, and an “NA” for the standard error estimate indicates that the specific value for the parameter was forced, as opposed to estimated. The penultimate column lists the log-likelihood value associated with the model, as computed through (13). Table 2 also lists Akaike’s Information Criterion (AIC)^72^ given by AIC = −2 log(max. likelihood) + 2*k*, where *k* is the number of parameters in the model. AIC is an information theoretic criterion that estimates the average Kullback-Liebler divergence of the estimated model from the true model; the term 2*k* should be interpreted as a penalty to encourage parsimonious models, with smaller AIC values indicating more favorable models.

**Table 2.**
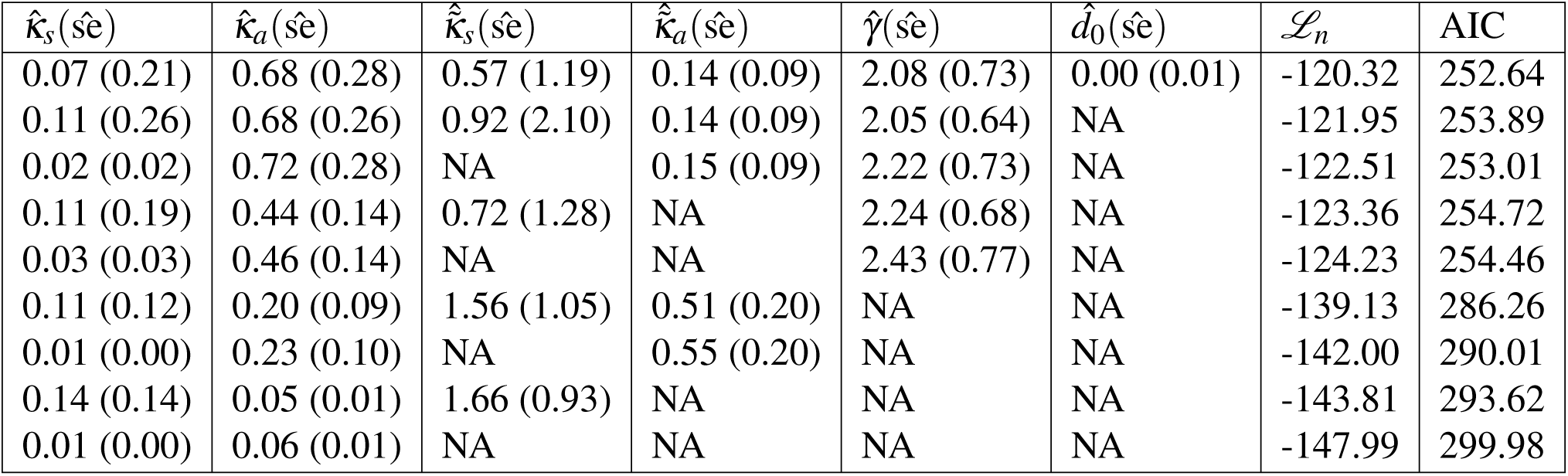
Model Estimation Output From Several Competing Models for MD-MBA-231.

Several observations about the results in Table 2 are salient, and lead to our final model recommendation. For example, the discounting parameter *γ* which connotes the ability of a tumor to recuperate after a sub-lethal encounter with the CAR T cell, is crucial for low AIC (and high likelihood) values as reflected in the first five entries of Table 2. Furthermore, estimates of *γ* remain stable with low standard error estimates, across the first five table entries. By contrast, the parameter *d*_0_ (connoting the baseline death rate) was found to be statistically insignificant. This is unsurprising because there were no observed cancer cell deaths in the absence of CAR T cells. Similarly, the parameter *κ̃*_*s*_ which is a measure of CAR T cell exhaustion, turns out to be statistically insignificant, as reflected in the first, second, and fourth entries of Table 2. The preceding discussion suggests the third and the fifth entries of Table 2 as the final competing models. Either of these models can serve as an adequate representation of the cell:cell dynamics, with the third entry being preferred due to having a slightly lower AIC value. The third entry in Table 2 is also interesting due to its inclusion of the *κ̃*_*a*_ parameter, providing evidence of the existence of adherent CAR T “exhaustion hypothesis.”

### Supplemental figures

**Figure S1.**
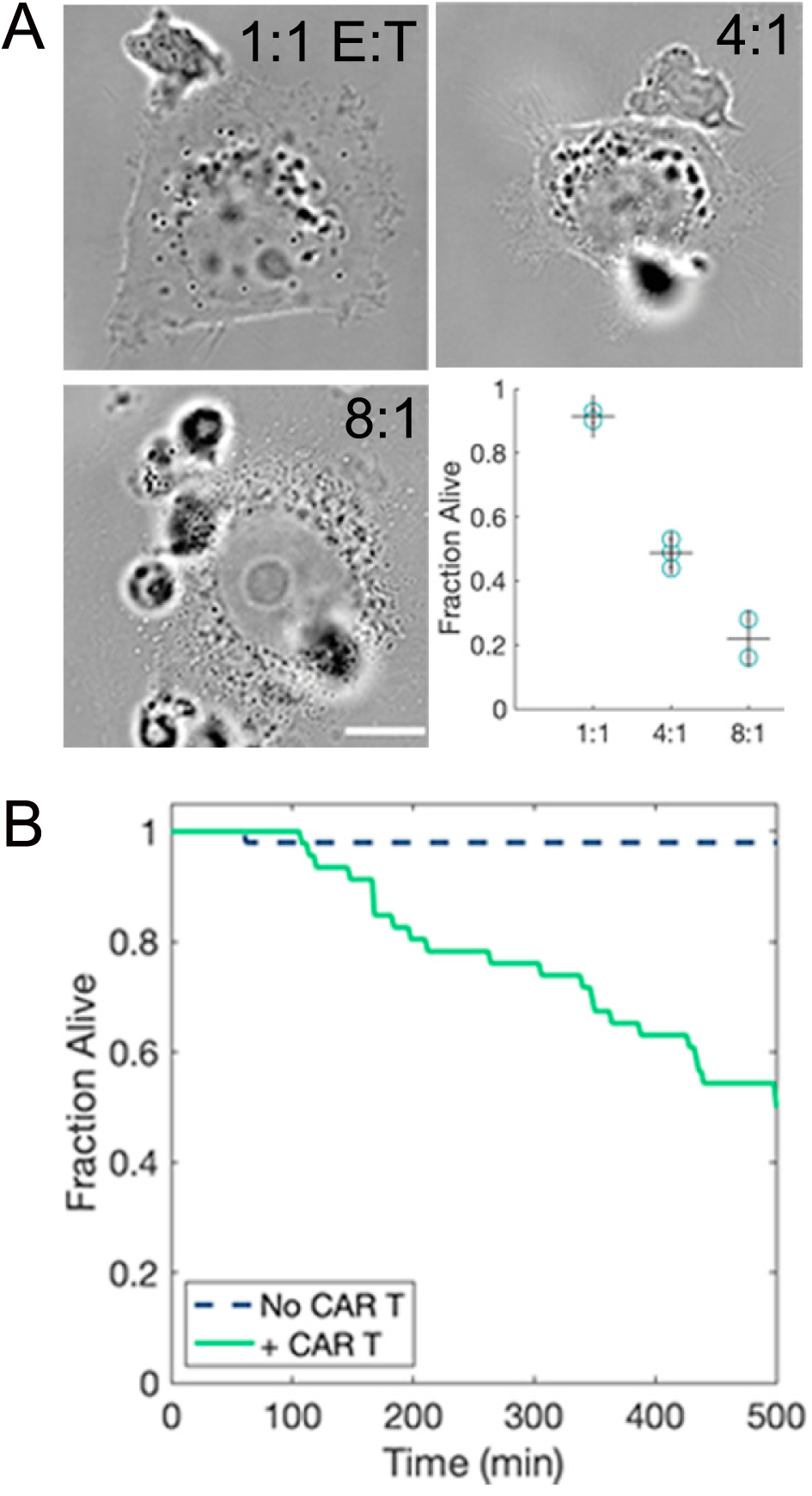
Killing outcomes increase with increased effector:target. A. Example diascopic images at varying E:T ratios and fraction of target cells alive after 16 hours (lower right). Data are plotted showing the biological replicates, mean, and standard deviation. Scale bar = 10 microns. B. Kaplan-Meier plot for MDA-MB-231 cells over 16 hours in the absence (dashed line) or presence (solid line) of CAR T cells at a 4:1 E:T ratio.

**Figure S2.**
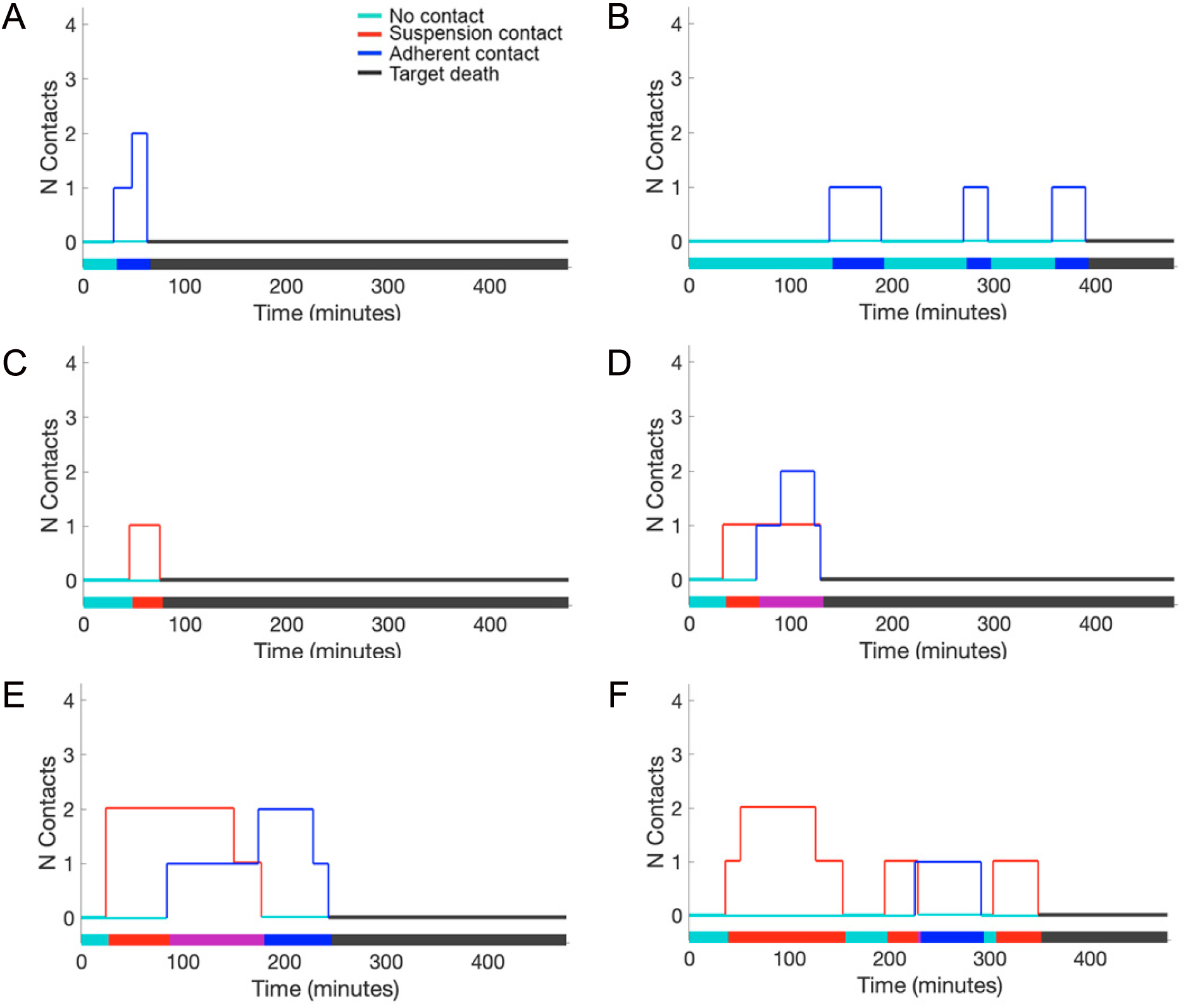
Additional example state traces for contact sequences that resulted in target cell death. CAR T cell:target contacts in the suspension (red) and adherent contexts (blue) are shown as a function of time. Target alive shown in teal and target dead indicated in gray. The overall history is shown as a projected timeline along the x-axis with the types of contact, where simultaneous adherent and suspension contacts are in purple, and the state of the target cell as a function of time.

**Figure S3.**
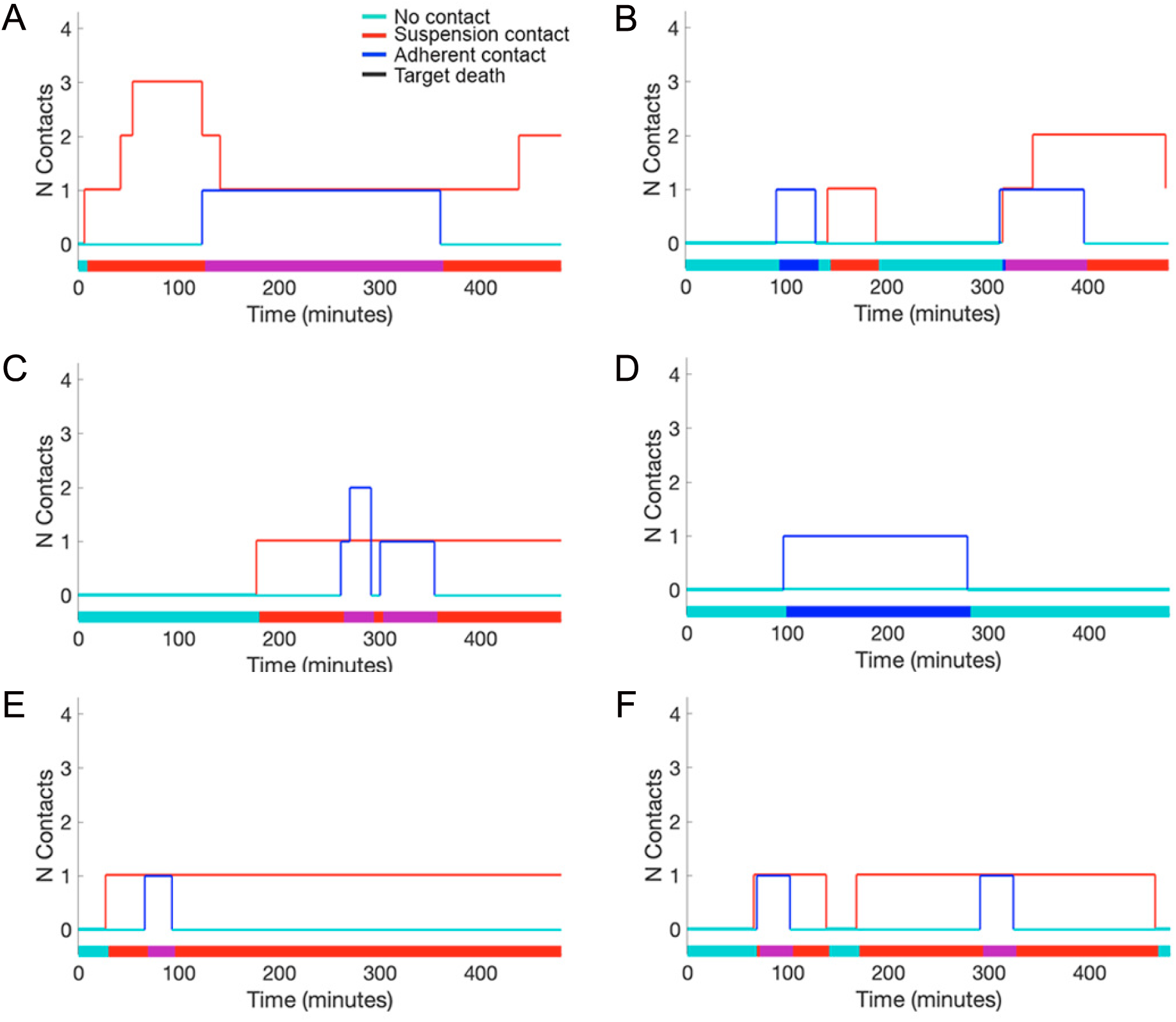
Additional example state traces that result in target cell survival. CAR T cell: target contacts in the suspension (red) and adherent contexts (blue) are shown as a function of time. Target alive shown in teal. The overall history is shown as a projected timeline along the x-axis with the types of contact, where simultaneous adherent and suspension contacts are in purple, and the state of the target cell as a function of time.

**Figure S4.**
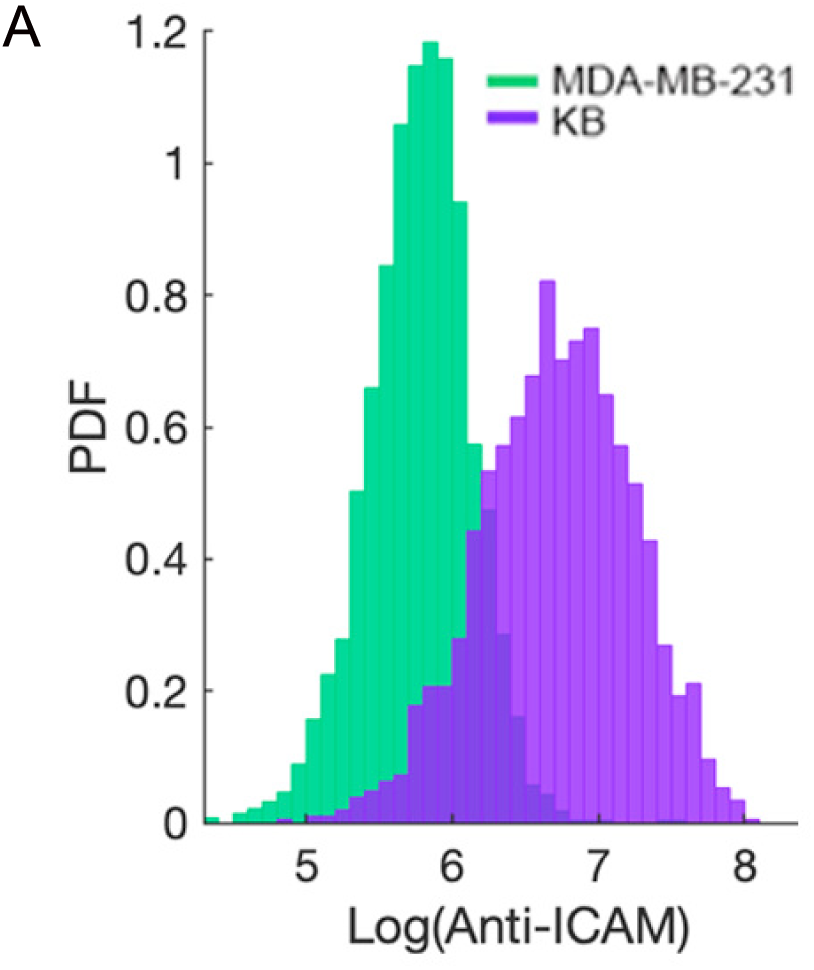
Flow cytometric detection of ICAM expression on MDA-MB-231 and KB cells.

**Figure S5.**
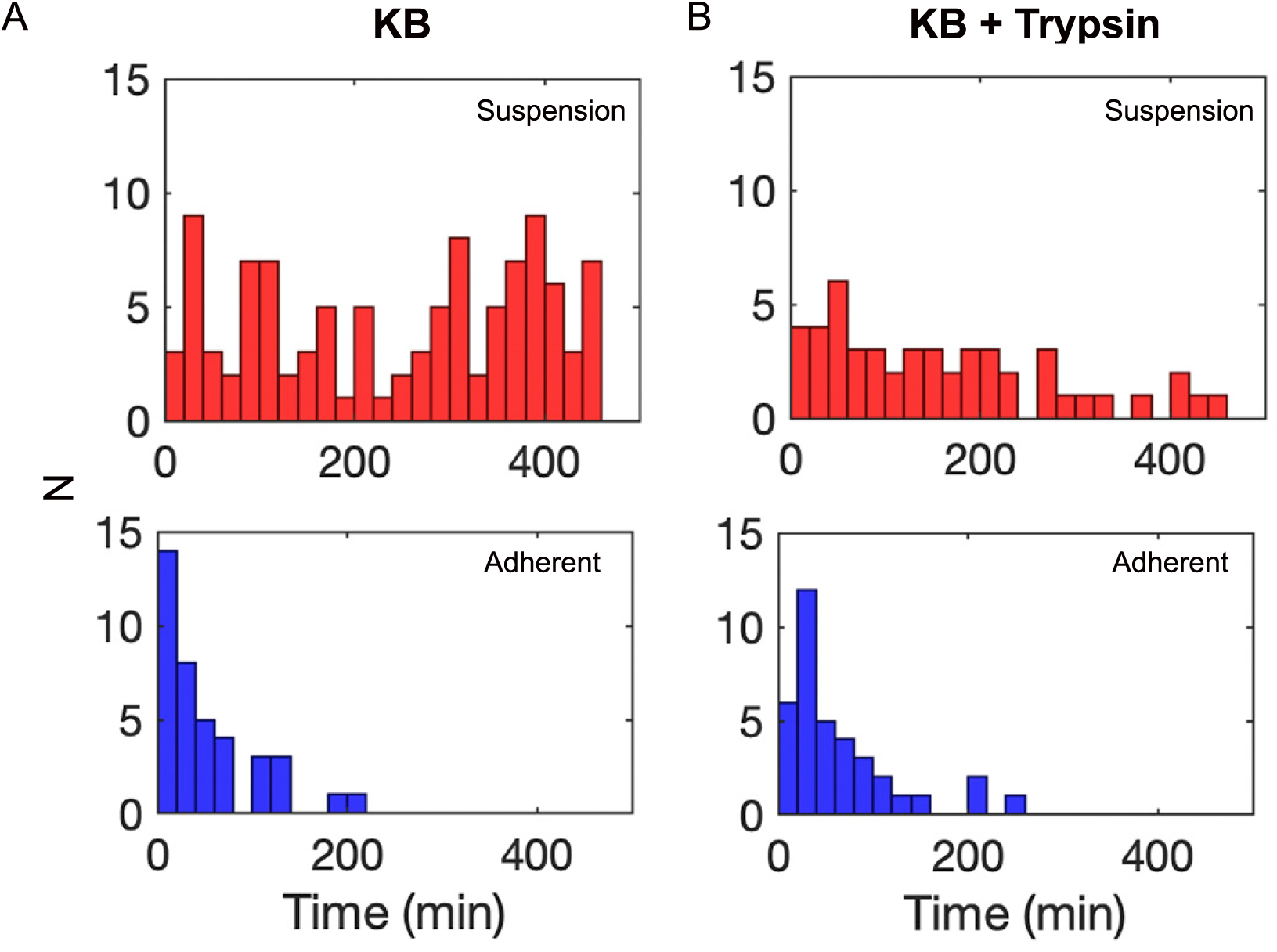
Interaction time distributions for the suspension (red; top row) and adherent (blue; bottom row) contact modes for KB cells (left column) and KB cells treated with trypsin (right column).

**Figure S6.**
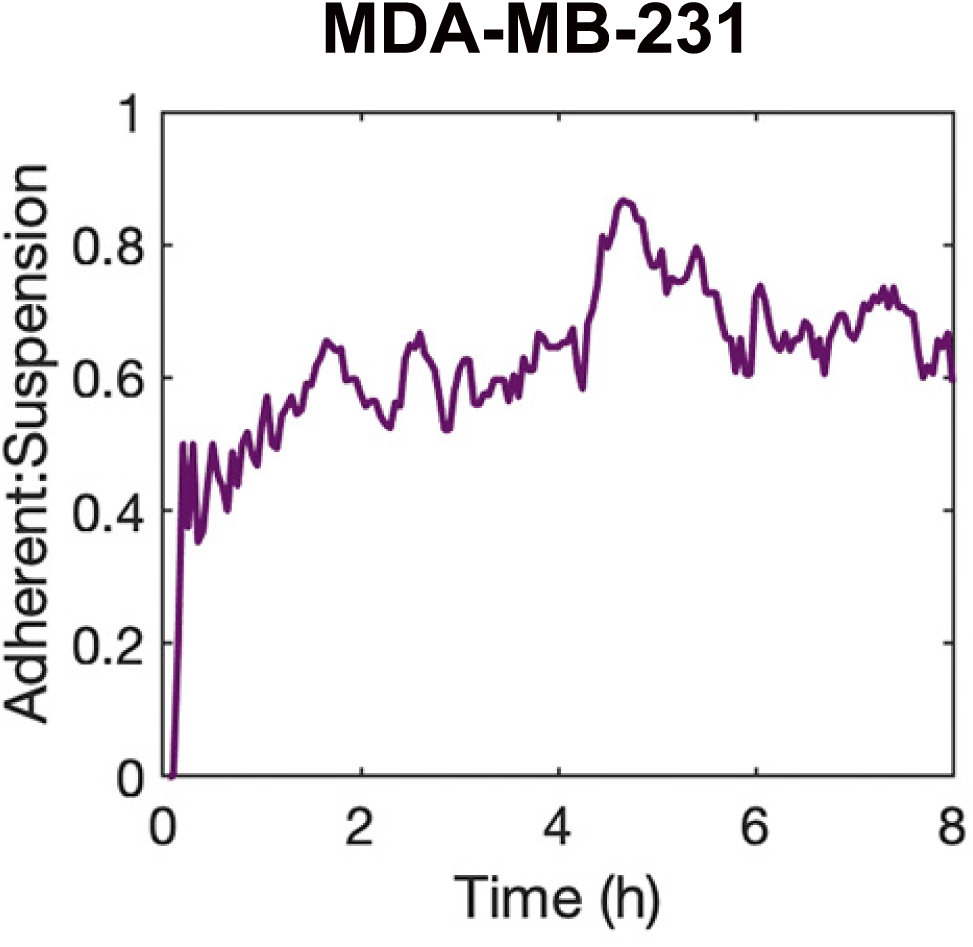
CAR T cell arrivals. Average number of suspension (red) and adherent (blue) contacts with MDA-MB-231 target cells as a function of time.

**Figure S7.**
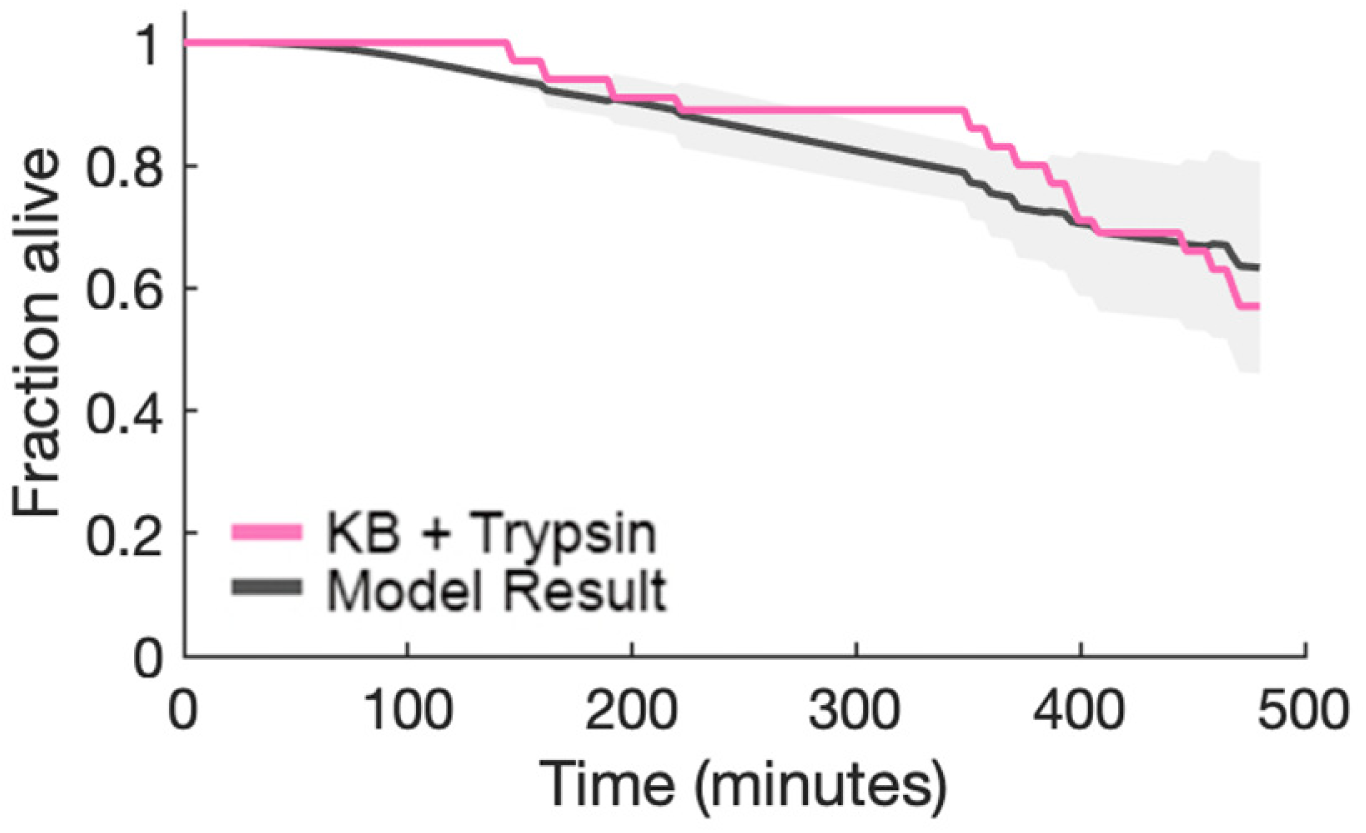
Model result conditioned on KB cells treated with trypsin. Kaplan-Meier plot for 40 simulations. The average killing curve is shown in gray with the upper and lower bound estimators shown as a light gray area around the mean. The empirical data for destruction of KB target cells treated with trypsin are plotted in pink.

### Supplemental movies

**Movie 1.** Example of CAR T cell contact with MDA-MB-231 target and resulting tumor cell death. Interaction of a single CAR T cell, loaded with fluorescent LysoTracker Deep Red (shown in magenta) interacting with a target MDA-MB-231 cell plated on glass substrate. The CAR T cell approaches and contacts the target, polarizes its cytotoxic payload, and destroys the target. Montage for these data shown in Fig. 1C.

**Movie 2.** Example of simultaneous imaging of the same CAR T cells and target cell in diascopic and IRM modes. Left: Diascopic imaging of a target cell in the presence of multiple CAR T cells. Right: Corresponding IRM images. The CAR T cell experiences contact in the adherent mode and burrows underneath the target cell. Montage for these data shown in Fig. 1E.

### Data availability

The data associated with this manuscript are available from the corresponding authors on request.

### Code availability

The code associated with this manuscript is available from the corresponding authors on request.

## Acknowledgments

We thank Tanvi Breinig for assistance in culturing primary T cells and all members of the Low-Nam team for their feedback. The authors acknowledge critical feedback from Geoffrey Graff (Purdue University). The work was supported by the Ralph W. and Grace M. Showalter Research Trust.

## Author contributions statement

S.S., R.P., and S.L.-N. designed the measurements, interpreted the data, developed and evaluated the model, and wrote the manuscript. S.S., K.S., and K.R-L. established the cell:cell killing assays and collected the data. S.S. curated the data and ran the models. S.Z., B.H., and P.L. provided the CAR T cells. All authors read and commented on the manuscript.

## Competing interests

The authors declare no competing interests.

## Notes

### Competing Interest Statement

The authors have declared no competing interest.

## References

1. Almåsbak, H., Aarvak, T., Vemuri, M. C. et al. Car t cell therapy: a game changer in cancer treatment. J. immunology research 2016 (2016).

2. Graham, C., Hewitson, R., Pagliuca, A. & Benjamin, R. Cancer immunotherapy with car-t cells– behold the future. *Clin*. Medicine 18, 324 (2018).

3. Davenport, A. J. et al. Car-t cells inflict sequential killing of multiple tumor target cells. Cancer immunology research 3, 483–494 (2015).

4. Benmebarek, M.-R. et al. Killing mechanisms of chimeric antigen receptor (car) t cells. Int. journal molecular sciences 20, 1283 (2019).

5. Weigelin, B. et al. Cytotoxic t cells are able to efficiently eliminate cancer cells by additive cytotoxicity. *Nat*. communications 12, 5217 (2021).

6. Liadi, I. et al. Individual motile cd4+ t cells can participate in efficient multikilling through conjugation to multiple tumor cells. Cancer immunology research 3, 473–482 (2015).

7. Halle, S. et al. In vivo killing capacity of cytotoxic t cells is limited and involves dynamic interactions and t cell cooperativity. Immunity 44, 233–245, 10.1016/j.immuni.2016.01.010 (2016).

8. Martinez, M. & Moon, E. K. Car t cells for solid tumors: new strategies for finding, infiltrating, and surviving in the tumor microenvironment. *Front*. immunology 10, 128 (2019).

9. Sterner, R. C. & Sterner, R. M. Car-t cell therapy: current limitations and potential strategies. Blood cancer journal 11, 69 (2021).

10. Shah, N. N. & Fry, T. J. Mechanisms of resistance to car t cell therapy. Nat. reviews Clin. Oncology 16, 372–385 (2019).

11. Nguyen, D. T., et al. Car t cell locomotion in solid tumor microenvironment, 10.3390/cells11121974 (2022).

12. Bonnet, V. et al. Cancer-on-a-chip model shows that the adenomatous polyposis coli mutation impairs t cell engagement and killing of cancer spheroids. 10.1073/pnas (2024).

13. Ronteix, G. et al. High resolution microfluidic assay and probabilistic modeling reveal cooperation between t cells in tumor killing. Nat. Commun. 13, 10.1038/s41467-022-30575-2 (2022).

14. Schnalzger, T. E. et al. 3d model for car-mediated cytotoxicity using patient-derived colorectal cancer organoids. The EMBO journal 38, e100928 (2019).

15. Ronteix, G. et al. High resolution microfluidic assay and probabilistic modeling reveal cooperation between t cells in tumor killing. *Nat*. communications 13, 3111 (2022).

16. Vasconcelos, Z. et al. Individual human cytotoxic t lymphocytes exhibit intraclonal heterogeneity during sustained killing. Cell Reports 11, 1474–1485 (2015).

17. Rothstein, T. L., Mage, M., Jones, G. & Mchugh, L. L. Cytotoxic t lymphocyte sequential kh,l g of immobilized allogeneic tumor target cells measured by time-lapse microcinematography (1978).

18. Perelson’, A. S., & Bell, G. I. Delivery of lethal hits by cytotoxic t lymphocytes in multicellular conjugates occurs sequentially but at random times’.

19. Salter, A. I. et al. Comparative analysis of tcr and car signaling informs car designs with superior antigen sensitivity and in vivo function. Tech. Rep. (2021).

20. Roybal, K. T. et al. Precision tumor recognition by t cells with combinatorial antigen-sensing circuits. Cell 164, 770–779, 10.1016/j.cell.2016.01.011 (2016).

21. Lim, W. A. & June, C. H. The principles of engineering immune cells to treat cancer, 10.1016/j.cell.2017.01.016 (2017).

22. Davenport, A., et al. Chimeric antigen receptor t cells form nonclassical and potent immune synapses driving rapid cytotoxicity. Proc. Natl. Acad. Sci. 115, E2068–E2076 (2018).

23. Xu, X. et al. Challenges and clinical strategies of car t-cell therapy for acute lymphoblastic leukemia: overview and developments. *Front*. immunology 11, 569117 (2021).

24. Liu, D. et al. The role of immunological synapse in predicting the efficacy of chimeric antigen receptor (car) immunotherapy. Cell communication signaling 18, 1–20 (2020).

25. Gudipati, V. et al. Inefficient car-proximal signaling blunts antigen sensitivity. Nat. Immunol. 21, 848–856, 10.1038/s41590-020-0719-0 (2020).

26. Xiong, Y., Libby, K. A. & Su, X. The physical landscape of car-t synapse, 10.1016/j.bpj.2023.09.004 (2024).

27. Dong, R. et al. Rewired signaling network in t cells expressing the chimeric antigen receptor (car). The EMBO J. 39, 1–14, 10.15252/embj.2020104730 (2020).

28. Yuan, D. J., Shi, L. & Kam, L. C. Biphasic response of t cell activation to substrate stiffness. Biomaterials 273, 120797 (2021).

29. Saitakis, M. et al. Different tcr-induced t lymphocyte responses are potentiated by stiffness with variable sensitivity. Elife 6, e23190 (2017).

30. O’Connor, R. S. et al. Substrate rigidity regulates human t cell activation and proliferation. The J. Immunol. 189, 1330–1339 (2012).

31. Chaudhuri, P. K., Wang, M. S., Black, C. T., Huse, M. & Kam, L. C. Modulating t cell activation using depth sensing topographic cues. Adv. Biosyst. 4, 10.1002/adbi.202000143 (2020).

32. Zhao, L. et al. T cell engineering for cancer immunotherapy by manipulating mechanosensitive force-bearing receptors, 10.3389/fbioe.2023.1220074 (2023).

33. Liu, Y. et al. Cell softness prevents cytolytic t-cell killing of tumor-repopulating cells. Cancer Res. 81, 476–488, 10.1158/0008-5472.CAN-20-2569 (2021).

34. Zhou, Y. et al. Cell softness renders cytotoxic t lymphocytes and t leukemic cells resistant to perforin-mediated killing. Nat. Commun. 15, 10.1038/s41467-024-45750-w (2024).

35. Martinez, M. & Moon, E. K. Car t cells for solid tumors: new strategies for finding, infiltrating, and surviving. Front. Immunol. 10, 128, 10.3389/fimmu.2019.00128 (2019).

36. Cazaux, M. et al. Single-cell imaging of car t cell activity in vivo reveals extensive functional and anatomical heterogeneity. J. Exp. Medicine 216, 1038–1049 (2019).

37. Halle, S. et al. In vivo killing capacity of cytotoxic t cells is limited and involves dynamic interactions and t cell cooperativity. Immunity 44, 233–245 (2016).

38. Wilkinson, D. J. Stochastic modelling for systems biology (Chapman and Hall/CRC, 2018).

39. Ross, S. M. Introduction to probability models (Academic press, 2014).

40. Wasserman, L. *All of statistics : a concise course in statistical inference*. Springer Texts in Statistics (Springer New York, New York, NY, 2004).

41. O’Quigley, J., et al. Survival Analysis (Springer, 2021).

42. Lu, Y. J. et al. Preclinical evaluation of bispecific adaptor molecule controlled folate receptor car-t cell therapy with special focus on pediatric malignancies. Front. oncology 9, 151 (2019).

43. Lu, Y. J. et al. Preclinical evaluation of bispecific adaptor molecule controlled folate receptor car-t cell therapy with special focus on pediatric malignancies. Front. Oncol. 9, 1–20, 10.3389/fonc.2019.00151 (2019).

44. Weigelin, B. et al. Cytotoxic t cells are able to efficiently eliminate cancer cells by additive cytotoxicity. Nat. Commun. 12, 10.1038/s41467-021-25282-3 (2021).

45. Ritter, A. T. et al. Actin depletion initiates events leading to granule secretion at the immunological synapse. Immunity 42, 864–876, 10.1016/j.immuni.2015.04.013 (2015).

46. Basu, R. et al. Cytotoxic t cells use mechanical force to potentiate target cell killing. Cell 165, 100–110, 10.1016/j.cell.2016.01.021 (2016).

47. Judokusumo, E., Tabdanov, E., Kumari, S., Dustin, M. L. & Kam, L. C. Mechanosensing in t lymphocyte activation. Biophys. J. 102, L5–L7, 10.1016/j.bpj.2011.12.011 (2012).

48. Tamzalit, F., et al. Interfacial actin protrusions mechanically enhance killing by cytotoxic t cells. Sci. Immunol. 4, 10.1126/sciimmunol.aav5445 (2019).

49. Blumenthal, D., Chandra, V., Avery, L. & Burkhardt, J. K. Mouse t cell priming is enhanced by maturation-dependent stiffening of the dendritic cell cortex. eLife 9, 1–44, 10.7554/eLife.55995 (2020).

50. Tello-Lafoz, M. et al. Cytotoxic lymphocytes target characteristic biophysical vulnerabilities in cancer. Immunity 54, 1037–1054.e7, 10.1016/j.immuni.2021.02.020 (2021).

51. Montalvo, M. J. et al. Decoding the mechanisms of chimeric antigen receptor (car) t cell-mediated killing of tumors: insights from granzyme and fas inhibition. Cell Death Dis. 15, 10.1038/s41419-024-06461-8 (2024).

52. Kumari, S., et al. Cytoskeletal tension actively sustains the migratory t-cell synaptic contact. The EMBO J. 39, 10.15252/embj.2019102783 (2020).

53. Jung, P., Zhou, X., Iden, S., Bischoff, M. & Qu, B. T cell stiffness is enhanced upon formation of immunological synapse. 10, 66643, 10.7554/eLife (2021).

54. Cazaux, M. et al. Single-cell imaging reveals car t cell heterogeneity. J. Exp. Medicine 216, 1038– 1049, 10.1084/jem.20182375 (2019).

55. Lopez, J. A. et al. Perforin forms transient pores on the target cell plasma membrane to facilitate rapid access of granzymes during killer cell attack. 10.1182/blood-2012-07 (2013).

56. Almåsbak, H., Aarvak, T., Vemuri, M. C. et al. Car t cell therapy: a game changer in cancer treatment. J. immunology research 2016 (2016).

57. Rudloff, M. W. et al. Hallmarks of cd8+ t cell dysfunction are established within hours of tumor antigen encounter before cell division. Nat. Immunol. 24, 1527–1539, 10.1038/s41590-023-01578-y (2023).

58. Anikeeva, N. et al. Efficient killing of tumor cells by car-t cells requires greater number of engaged cars than tcrs. J. Biol. Chem. 297, 10.1016/J.JBC.2021.101033 (2021).

59. Boissonnas, A. et al. Cd8+ tumor-infiltrating t cells are trapped in the tumor-dendritic cell network. Neoplasia (United States) 15, 85–94, 10.1593/neo.121572 (2013).

60. Adu-Berchie, K. et al. Generation of functionally distinct t-cell populations by altering the viscoelasticity of their extracellular matrix. *Nat*. Biomed. Eng. 7, 1374–1391, 10.1038/s41551-023-01052-y (2023).

61. Park, S. et al. Immunoengineering can overcome the glycocalyx armour of cancer cells. Nat. Mater. 23, 429–438, 10.1038/s41563-024-01808-0 (2024).

62. Hou, A. J., Chen, L. C. & Chen, Y. Y. Navigating car-t cells through the solid-tumour microenvironment, 10.1038/s41573-021-00189-2 (2021).

63. de Jesus, M., et al. Single-cell topographical profiling of the immune synapse reveals a biomechanical signature of cytotoxicity. Sci. Immunol. 9, 10.1126/sciimmunol.adj2898 (2024).

64. Govendir, M. A. et al. T cell cytoskeletal forces shape synapse topography for targeted lysis via membrane curvature bias of perforin. Dev. Cell 57, 2237–2247.e8, 10.1016/j.devcel.2022.08.012 (2022).

65. Ritter, A. T. et al. Cortical actin recovery at the immunological synapse leads to termination of lytic granule secretion in cytotoxic t lymphocytes. Proc. Natl. Acad. Sci. United States Am. 114, E6585–E6594, 10.1073/pnas.1710751114 (2017).

66. Chitirala, P. et al. Studying the biology of cytotoxic t lymphocytes in vivo with a fluorescent granzyme b-mtfp knock-in mouse. eLife 9, 1–19, 10.7554/eLife.58065 (2020).

67. Breart, B., Lemaître, F., Celli, S. & Bousso, P. Two-photon imaging of intratumoral cd8+ t cell cytotoxic activity during adoptive t cell therapy in mice. J. Clin. Investig. 118, 1390–1397, 10.1172/JCI34388 (2008).

68. Torro, R. et al. Celldetective: an ai-enhanced image analysis tool for unraveling dynamic cell interactions, 10.7554/eLife.105302.1 (2025).

69. Ross, S. M. Introduction to probability models (Academic press, 2014).

70. Wasserman, L. All of statistics : a concise course in statistical inference (Springer New York, 2004).

71. Lehmann, E. L. Elements of large-sample theory (Springer, 1999).

72. Hurvice, C. M. & Tsai, C. L. Regression and time series model selection in small samples. Biometrika 76, 297–307 (1989).

